# Novel tools to quantify total, phospho-Ser129 and aggregated alpha-synuclein in the mouse brain

**DOI:** 10.1101/2024.06.12.598745

**Authors:** Benjamin Guy Trist, Courtney Jade Wright, Alejandra Rangel, Louise Cottle, Asheeta Prasad, Nanna Møller Jensen, Hjalte Gram, Nicolas Dzamko, Poul Henning Jensen, Deniz Kirik

## Abstract

Assays for quantifying aggregated and phosphorylated (S129) human α-synuclein protein are widely used to evaluate pathological burden in patients suffering from synucleinopathy disorders. Many of these assays, however, do not cross-react with mouse α-synuclein or exhibit poor sensitivity for this target, which is problematic considering the preponderance of mouse models at the forefront of pre-clinical α-synuclein research. In this project, we addressed this unmet need by reformulating two existing AlphaLISA^®^ SureFire^®^ Ultra^TM^ total and pS129 α-synuclein assay kits to yield robust and ultrasensitive (LLoQ ≤0.5pg/mL) quantification of mouse and human wild-type and pS129 α-synuclein protein. We then employed these assays, together with the BioLegend α-synuclein aggregate ELISA, to assess the relationship between α-synuclein S129 phosphorylation and aggregation in different mouse brain tissue preparations. Overall, we highlight the compatibility of these new immunoassays with rodent models and demonstrate their potential to advance knowledge surrounding α-synuclein phosphorylation and aggregation in synucleinopathies.

## Introduction

Synucleinopathies are a group of clinically heterogenous neurodegenerative disorders characterized by a unifying pathological feature: the abnormal deposition of Lewy pathology within neuronal and/or glial cell bodies and processes^1^. Within this broader classification the main subtypes are Parkinson’s disease, representing the commonly seen clinical presentation, and two rare variants: dementia with Lewy bodies (DLB) and multiple system atrophy (MSA), the latter manifesting as two distinct subgroups exhibiting predominant symptoms of either cerebellar ataxia or parkinsonism. There are currently no disease modifying treatments capable of slowing or halting cell death in any of these disorders, however the abundance of Lewy pathology in brain regions undergoing significant functional decline and degeneration has made it a focal point for the development of such interventions.

The structural morphology and composition of Lewy pathology is highly heterogeneous across synucleinopathies^2^, as well as within and between different brain regions of each disorder^3,4^, complicating attempts to understand mechanisms driving its formation. Lewy pathology broadly comprises spherical Lewy bodies (LBs) and pale bodies, as well as filamentous Lewy neurites (LNs), which are composed of dysmorphic organellar components, lipid membranes^5^ and many hundreds of proteins^6^. Despite such incredible variability in composition, α-synuclein protein is invariably enriched in all forms of Lewy pathology and is suggested to play a key role in its propagation^3^. The leading hypothesis proposes that intraneuronal α-synuclein misfolds, oligomerizes and transforms into β-sheet-rich amyloid fibrils^7^, which form the foundation of neuronal Lewy pathology. Furthermore, intermediate misfolded forms of α-synuclein are thought to be transferred to neighbouring neuronal and glial cells upon their release^8,9^. The molecular pathways governing α-synuclein misfolding, aggregation, fibrillization, transmission and Lewy pathology formation, however, remain poorly understood.

Post-translational modifications (PTMs) are chemical modifications to amino acid residue side chains of a protein that impact biophysical properties such as surface charge, hydropathy, size and conformation, which can in turn influence protein misfolding, aggregation, subcellular localization, activity and degradation. A large number of PTMs are reported for α-synuclein protein – phosphorylation, ubiquitination, nitration, acetylation, truncation, SUMOylation, glutathionylation and glycosylation^10^, with phosphorylation of serine residue 129 (pS129) attracting particular interest due to its high abundance within Lewy pathology^11^. While this has led to the development of several robust assays capable of quantifying human wild-type (WT) and pS129 α-synuclein in brain tissue, blood and cerebrospinal fluid^11-16^, assay cross reactivity for mouse WT and pS129 α-synuclein is poor^12,16^. This is problematic given the preponderance and implementation of mouse models in research examining the physiological and pathological roles of α-synuclein in Lewy pathology, and the potential utilisation of these models in therapeutic development.

To address this critical unmet need, we worked with the manufacturers of the AlphaLISA^®^ SureFire^®^ Ultra^TM^ Total (PerkinElmer^®^, #ALSU-TASYN) and Phospho-α-Synuclein (Ser129; PerkinElmer^®^, #ALSU-PASYN) Detection Kits to adapt their formulations and ensure compatibility with both human and mouse α-synuclein. Comprehensive characterization of these reformulated assays demonstrates that they are capable of robust and highly sensitive (∼0.1-0.5 pg/mL) quantification of both human and mouse WT and pS129 α-synuclein *in vitro,* in complex cell lysates, as well as brain tissue extracts. Furthermore, we complemented these assays with the aggregated α-synuclein ELISA assay from BioLegend^®^ to provide the first demonstration of alterations to the total abundance, phosphorylation, aggregation, and subcellular compartmentalization of mouse α-synuclein within the brains of WT mice following inoculation with WT mouse α-synuclein pre-formed fibrils (PFFs).

## Results

### Selection and validation of new antibody pairs for reformulated SureFire Ultra pS129 and total α-synuclein assays using purified protein standards

Prior to this project, we discovered that the existing SureFire Ultra pS129 α-synuclein assay (#ALSU-PASYN-A, PerkinElmer) did not recognize human pS129 α-synuclein, while the existing total α-synuclein assay (#ALSU-TASYN-A, PerkinElmer) was unable to detect mouse pS129 or mouse WT α-synuclein (**Supplementary Table 1**). We therefore aimed to improve cross species detection in both of these assays by screening 5 new formulations of each assay against purified human and mouse pS129 and WT α-synuclein recombinant protein standards. Each new formulation contained different immunocapture antibody pairings, which were chosen based on their cross-reactivity for mouse and human α-synuclein isoforms in previous immunocapture or immunostaining experiments conducted by our group and others^11,12,17^. Accordingly, both species of pS129 α-synuclein were robustly detected above background signal produced from assay lysis buffer by 3 new pS129 α-synuclein assay formulations (**Figure 1a,b**) and 1 new total α-synuclein assay formulation (**Figure 1c,d**). This same total assay formulation was also the only one to detect both human and mouse WT α-synuclein (**Figure 1e,f**). Neither species of WT α-synuclein were detected by any pS129 assay formulation (**Figure 1g,h**).

**Figure 1.**
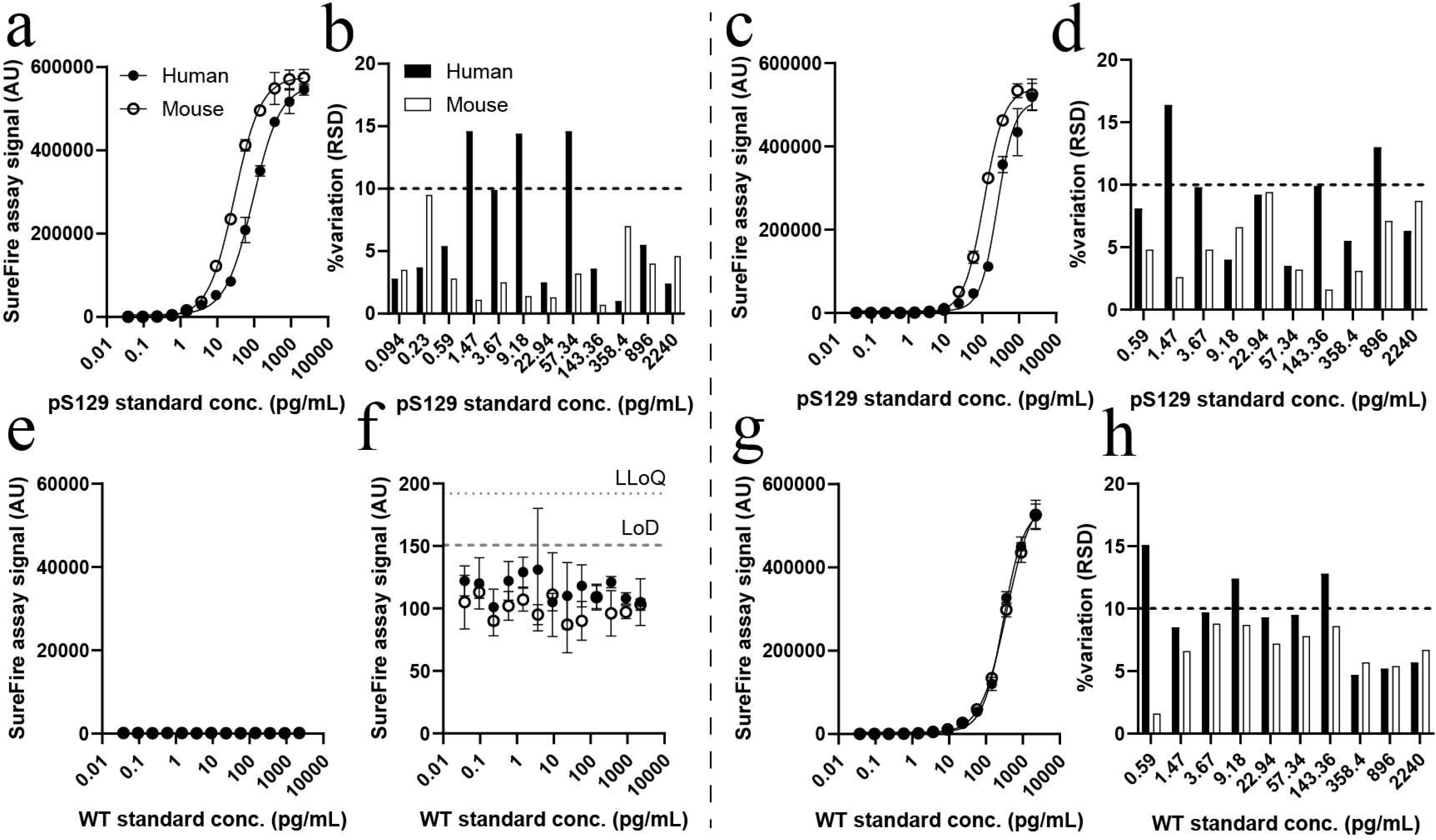
Characterization of new pS129 and total α-synuclein SureFire Ultra assays using purified protein standards. Five new formulations of the pS129 (left) and total (right) α-synuclein assay were characterized using purified human (Hu) and mouse (Ms) pS129 (**a-d**) and wild-type (WT; **e-h**) α-synuclein standards, with data presented above representing only the most sensitive formulation of each assay. Associated data for all tested formulations are contained within **Supplementary Tables 2-3**. Inter-assay variation (RSD) was calculated from three individual standard curves constructed and measured on separate days (**b, d, h**). The limit of detection (LoD) and lower limit of quantification (LLoQ) were defined as 3 and 6 standard deviations above the mean of the blank, respectively. Data in panels **a, c, e, g** represent mean ± standard deviation, while % variation data in panels **b, d, h** are only presented for standard concentrations detected above the LLoQ. AU, arbitrary units.

The most sensitive formulation of each assay yielded 5-to-20-fold improvements in sensitivity compared with previous SureFire Ultra α-synuclein assay formulations, and were also 10-to-20-fold more sensitive than alternative α-synuclein assays^12,18,19^. The most sensitive new pS129 assay exhibited a limit of detection (LoD) between 0.02-0.04 pg/mL and a lower limit of quantification (LLoQ) between 0.04-0.08 pg/mL for both human and mouse pS129 α-synuclein species. Its linear dynamic range was 5-fold greater than alternative assays (0.1 to 143.4 pg/mL; **Supplementary Figure 2**)^12,18,19^ and the average inter-assay variability was 5% (range 0.7-14.6%). Similarly, the new total assay formulation exhibited a LoD between 0.04-0.09 pg/mL and a LLoQ between 0.1-0.5 pg/mL for all standards. Its linear dynamic range was 10-fold greater than alternative assays^12,18,19^, ranging from 0.2-896 pg/mL for all standards except mouse pS129 α-synuclein (0.2-358 pg/mL; **Supplementary Figure 2**), with an average inter-assay variability of 8.0% (range 1.6-15.0%). Complete assay characterization data for purified protein standards is provided in **Supplementary Tables 1-3**.

### Assessment of reformulated assay sensitivity and selectivity in complex biological matrices

Screening new assay formulations *in vitro* informed on the compatibility of new antibody pairs and their selectivity for mouse and human α-synuclein isoforms in isolation, however these parameters can differ in complex biological matrices if target antigen conformations vary due to interactions with other cellular components. We cross-validated the *in vitro* sensitivity and selectivity of new assay formulations for both species of WT and pS129 α-synuclein in complex biological matrices by applying the most sensitive formulation of each assay to WT mouse brain tissue and HEK293 cell extracts. These assays will subsequently be referred to as the reformulated total and pS129 α-synuclein assays, which are now commercially available through Revvity (total, #ALSU-TASYN-B; pS129, #ALSU-PASYN-B). While pS129 α-synuclein is abundant in the WT mouse brain, it exists in comparatively low abundance in WT HEK293 cells^12^, hindering assessment of pS129 α-synuclein assay performance under baseline conditions in this cell line. To address this challenge, we transfected WT HEK293 cells with a vector construct expressing polo-like kinase 3 (PLK3) under a human CMV promoter^20^, dramatically increasing phosphorylation of α-synuclein at the S129 residue. Mirroring *in vitro* data, the reformulated pS129 α-synuclein assay subsequently robustly detected mouse and human pS129 α-synuclein in WT mouse brain and PLK3-transfected HEK293 cell extracts (**Figure 2a,b**), while the reformulated total α-synuclein assay detected mouse and human α-synuclein in WT mouse brain and HEK293 cell extracts (**Figure 2c,d**). Both assays again exhibited low inter-assay variability in these matrices (average 5.0%, range 0.8-19.6%).

**Figure 2.**
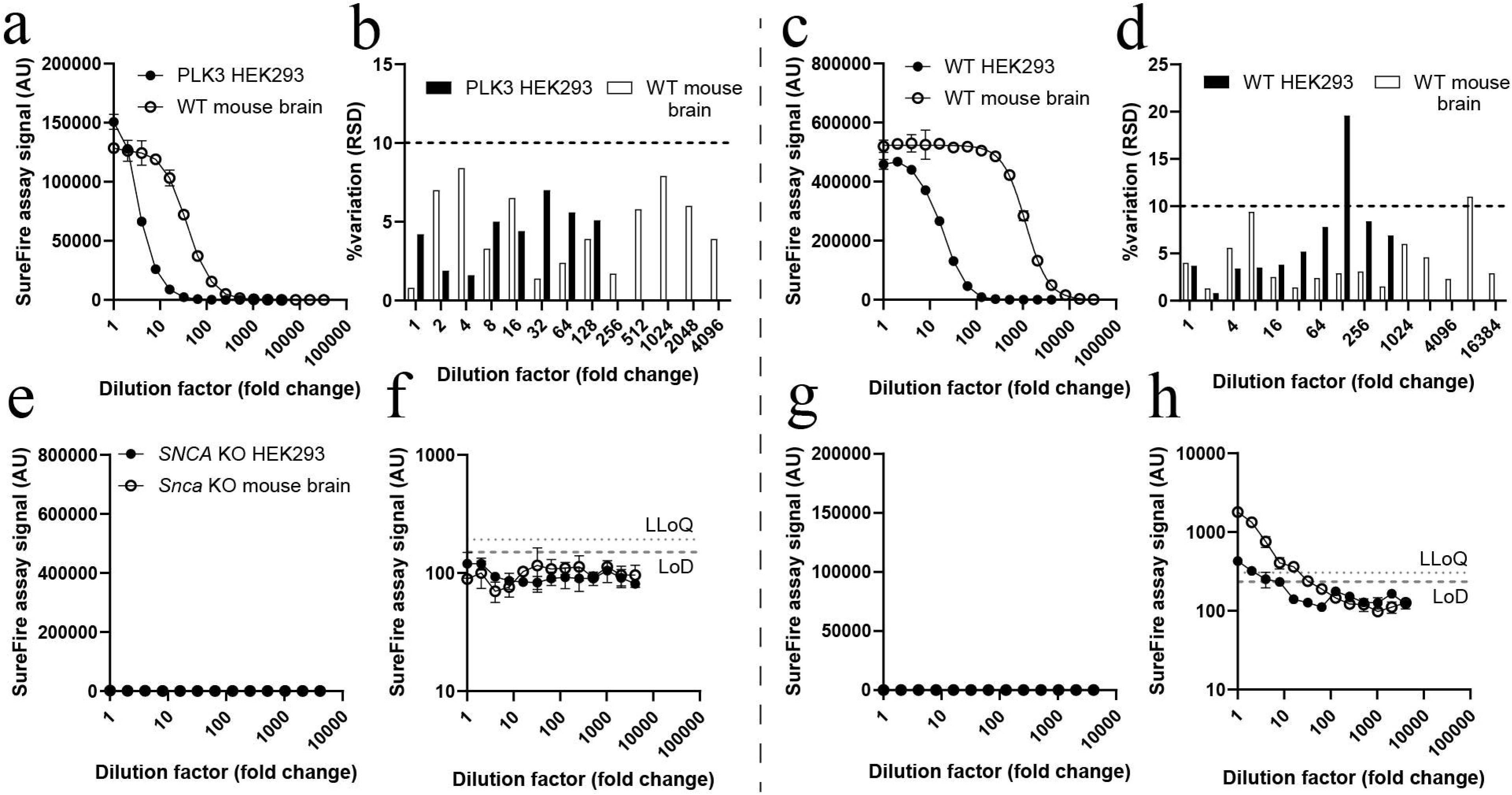
Characterization of new pS129 and total α-synuclein SureFire Ultra assays using mouse brain tissue extracts and HEK293 cell lysates. The most sensitive new pS129 α-synuclein assay (left) was characterized using wild-type (WT) mouse brain tissue extracts and PLK3-transfected HEK293 cell lysates (**a, b**), while testing of the most sensitive total α-synuclein assay (right) utilised WT mouse brain tissue extracts and WT HEK293 cell lysates (**c, d**). Extracts and lysates were serially-diluted using assay buffer. Associated assay data for all formulations tested are contained within **Supplementary Table 4**. Inter-assay variation (RSD) data presented in panels **b** and **d** were calculated from three individual dilution curves constructed and measured on separate days, with data only shown for extract/lysate dilutions above the lower limit of quantification (LLoQ). Phospho-S129 (**e, f**) and total (**g, h**) assay specificity was also assessed using *SNCA* knock-out (KO) HEK293 cell lysates and *Snca* KO mouse brain tissue extracts. The limit of detection (LoD) and LLoQ were defined as described in Figure 1. Data in panels **a, b, g, h** represent mean ± standard deviation. AU, arbitrary units.

To assess non-specific binding of assay reagents to other matrix components, we applied our reformulated pS129 and total α-synuclein assays to *Snca* knock-out (KO) mouse brain tissue extracts and *SNCA* KO HEK293 cell lysates diluted serially with assay lysis buffer. We did not detect any signal above the LLoQ for any dilution of either KO matrix using the pS129 assay, suggesting high specificity of this assay for pS129 α-synuclein (**Figure 2e,f**). Application of the total assay to both KO matrices produced a very small signal (<0.3-0.4% of max assay signal for corresponding WT matrices) above the LLoQ in the most concentrated sample dilutions tested (**Figure 2g,h**), suggesting that one or both antibodies within this formulation possess a very low affinity for one or more components of both matrices aside from α-synuclein. We determined that the minimum required dilution (MRD) to successfully ameliorate these background signals was approximately 24-fold in *Snca* KO mouse brain tissue extracts and <8-fold in *SNCA* KO HEK293 cell lysates. Importantly, both of these MRDs are lower in magnitude than the dilution factors required to reach the linear dynamic range in corresponding WT mouse brain (256-fold) and HEK293 cell (>8-fold) extracts, making it highly unlikely that these minor matrix effects distort true positive assay signals. Complete assay characterization data for mouse brain extracts and HEK293 cell lysates is provided in **Supplementary Table 4**.

### Evaluation of matrix effects and their impact on reformulated assay performance

A key step in validating any ligand-binding assay is evaluating whether the immunoaffinity characteristics of the assay differ when performed in calibrator matrix (assay lysis buffer) compared with sample matrices (mouse brain tissue or HEK293 cell extracts). Differences in these characteristics between matrices, known as “matrix effects”, commonly arise from specific or non-specific interactions between matrix components and capture/detection reagents or the target antigen itself, which can diminish assay accuracy and sensitivity. We conducted parallelism and spike-in experiments to confirm the suitability of our calibrator matrix (assay lysis buffer) for measuring endogenous α-synuclein in mouse brain tissue and human cell extracts.

In parallelism experiments, WT and α-synuclein KO mouse brain tissue extracts were first diluted 10-fold and 100-fold, constituting dilution factors above and below the MRD (24-fold) previously determined to ameliorate non-specific assay signals in *Snca* KO mouse brain extracts (**Figure 2h)**. Pre-diluted WT matrices were then serially diluted using either assay lysis buffer or the corresponding dilution of α-synuclein KO matrix (**Figure 3a**). While serial dilutions in assay lysis buffer progressively reduced the concentration of all components of the study matrix, dilutions made using corresponding KO matrices ensured all components remain constant throughout the sample-dilution response curve except α-synuclein, preserving any matrix effects throughout the dilution series. If matrix effects are indeed absent beyond the MRD, the sample-dilution response curves for WT matrix diluted in assay lysis buffer and KO matrix should coincide. We observed significant sample-dilution response curve shifts in 10-fold-diluted WT mouse brain extracts diluted in KO matrix compared with assay buffer for both reformulated assays (**Figure 3b,c**), which disappeared when extracts were diluted 100-fold (**Figure 3d,e**). These distortions were markedly different from false-positive signals observed in neat α-synuclein KO matrices using the total α-synuclein assay (**Figure 2h**), with sample-dilution response curve shifts demonstrating a dampening of assay signal in 10-fold-diluted matrices compared with 100-fold-diluted matrices. Parallelism experiments were also performed in WT and *SNCA* KO HEK293 cell extracts, revealing a lower magnitude matrix effect that was ameliorated by diluting extracts ≥20-fold (**Supplementary Figure 3**).

**Figure 3.**
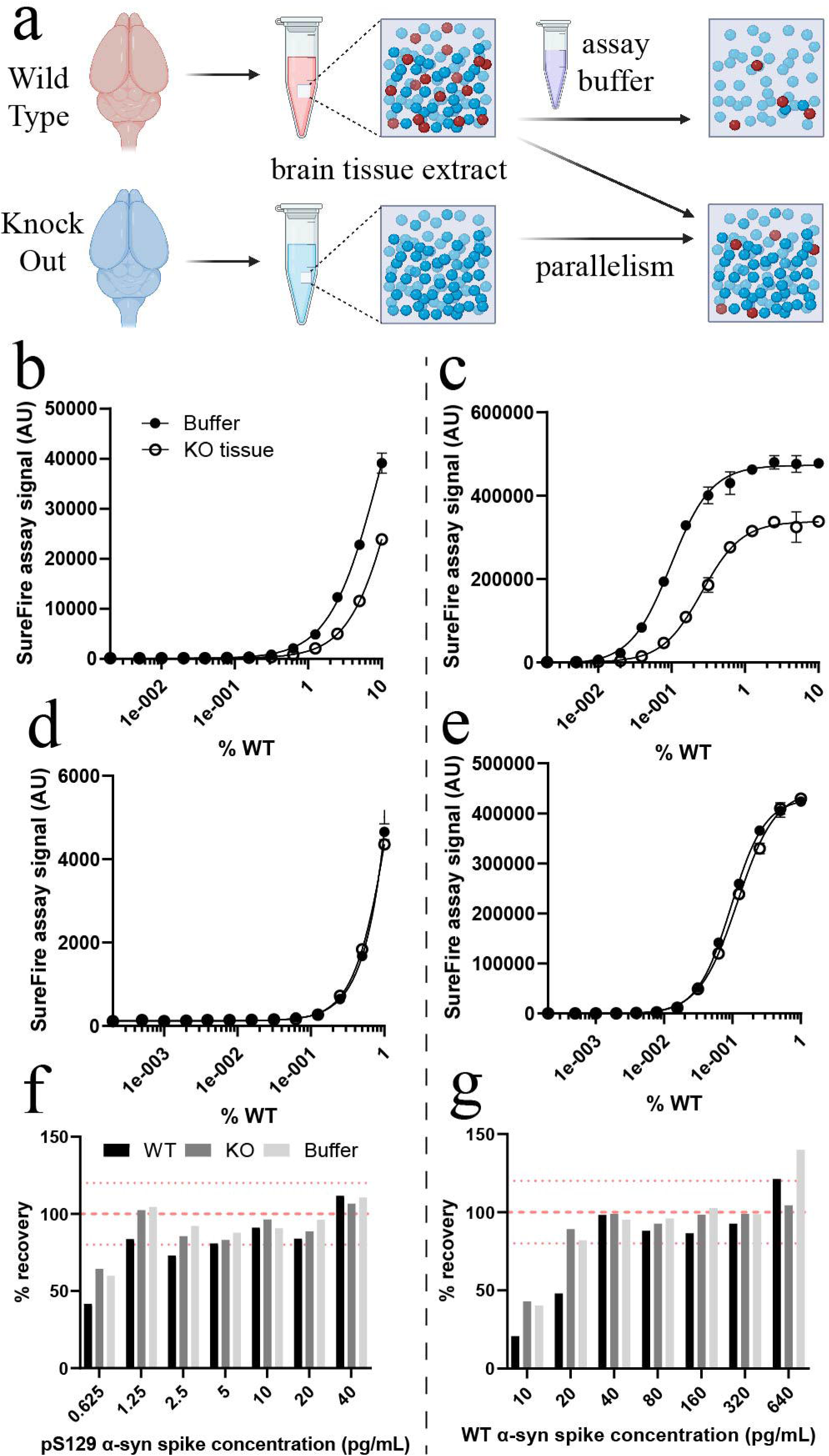
Evaluation of matrix effects and their impact on SureFire Ultra assay performance. The magnitude of matrix effects produced by mouse brain tissue was first evaluated in new assay formulations using parallelism experiments (**a**). Matrix effects were measured for the pS129 (left) and total (right) assay using wild-type (WT) and *Snca* knock-out (KO) mouse brain tissue extracts that had both been diluted 10-fold (**b, c**) and 100-fold (**d, e**). Phospho-S129 (**f**) and WT (**g**) mouse α-synuclein were also spiked into WT and *Snca* KO mouse brain tissue extracts diluted 2000-fold and 100-fold, respectively, as well as assay buffer (AB), and spike recovery assessed using standard curves generated for purified mouse WT and pS129 α-synuclein. Wild-type α-synuclein was spiked into samples at 10-640pg/mL and measured using the new total α-synuclein assay, while pS129 α-synuclein was spiked in at 0.625-40pg/mL and measured using the new pS129 α-synuclein assay. Dotted red lines in panels **f** and **g** represent 80% and 120% spike recovery, while dashed red lines in these panels represent 100% spike recovery. Data in panels **b-e** represent mean ± standard deviation. Associated data are contained within **Supplementary Table 5-7**. AU, arbitrary units.

For spike-in experiments, WT and α-synuclein KO mouse brain extracts were inoculated with known quantities of mouse WT and pS129 α-synuclein, before spike concentrations measured to assess the accuracy of α-synuclein quantification in these samples. Knock-out mouse brain extracts were diluted 100-fold to prevent matrix effects, while WT mouse brain extracts were further diluted to 2000-fold to ensure their α-synuclein concentration was on the lower end of the assays’ linear dynamic range. WT α-synuclein spikes were measured using the total assay and pS129 α-synuclein spikes measured using the pS129 assay. We observed proportionate increases in assay signal with increasing spike concentrations for pS129 and total assays, which did not differ significantly between matrices. Spike recovery from assay lysis buffer and mouse brain matrices averaged 80-92% for the pS129 assay (**Figure 3f**) and 79-94% for the total assay (**Figure 3g**), with notably lower spike recovery observed at lower spike concentrations. Spike recovery was especially poor for low spike concentrations made in wild-type mouse brain matrix, which we attribute to higher assay background signals produced by endogenous α-synuclein in this matrix. Complete assay characterization data for parallelism and spike-in experiments is provided in **Supplementary Tables 5-7**.

### Intrastriatal PFF-inoculation differentially alters α-synuclein abundance and phosphorylation across the WT mouse brain

The α-synuclein pre-formed fibril (PFF) model has become a widely used animal model of Parkinson’s disease, involving the central or peripheral injection of recombinant α-synuclein protein fibrils^21^ to trigger widespread α-synuclein phosphorylation and aggregation throughout the brain. However, tools to accurately quantify the concentration of these abnormal species of α-synuclein in the rodent brain are lacking. To determine whether our reformulated α-synuclein assays address this gap, we first performed bilateral striatal injections of mouse α-synuclein PFFs (5 mg/hemisphere) in WT mice to induce significant synucleinopathy, with corresponding PBS-injected mice (sham) serving as controls. Immunohistochemical profiling of pS129 α-synuclein was then conducted in fixed brain tissues from PFF (**Figure 4a,b**) and sham (**Supplementary Figure 4**) mice 3 months post-injection, and qualitative observations made to identify regions exhibiting high, moderate or no pS129 α-synuclein pathology. Cortical and limbic regions such as the motor cortex (MC), anterior cingulate cortex (ACC), somatosensory cortex (SSC) and amygdala (AMG) exhibited a high pathological burden in mouse α-synuclein PFF-treated mice, with other regions of these same brains such as the olfactory bulb (OLF), striatum (STR), hippocampus (HIP) and ventral midbrain (VMB) only possessing moderate pS129 α-synuclein pathology. All sham-treated mouse brain regions, as well as select regions of PFF-treated mice (cerebellum, CB; dorso-medial midbrain, DMB), showed minimal or no pathology. Following histological profiling, we applied both reformulated SureFire Ultra α-synuclein assays to whole tissue extracts from these same regions of the sham and PFF mouse brain to provide the first quantitative measurements of pS129 and total α-synuclein across the mouse brain following PFF inoculation.

**Figure 4.**
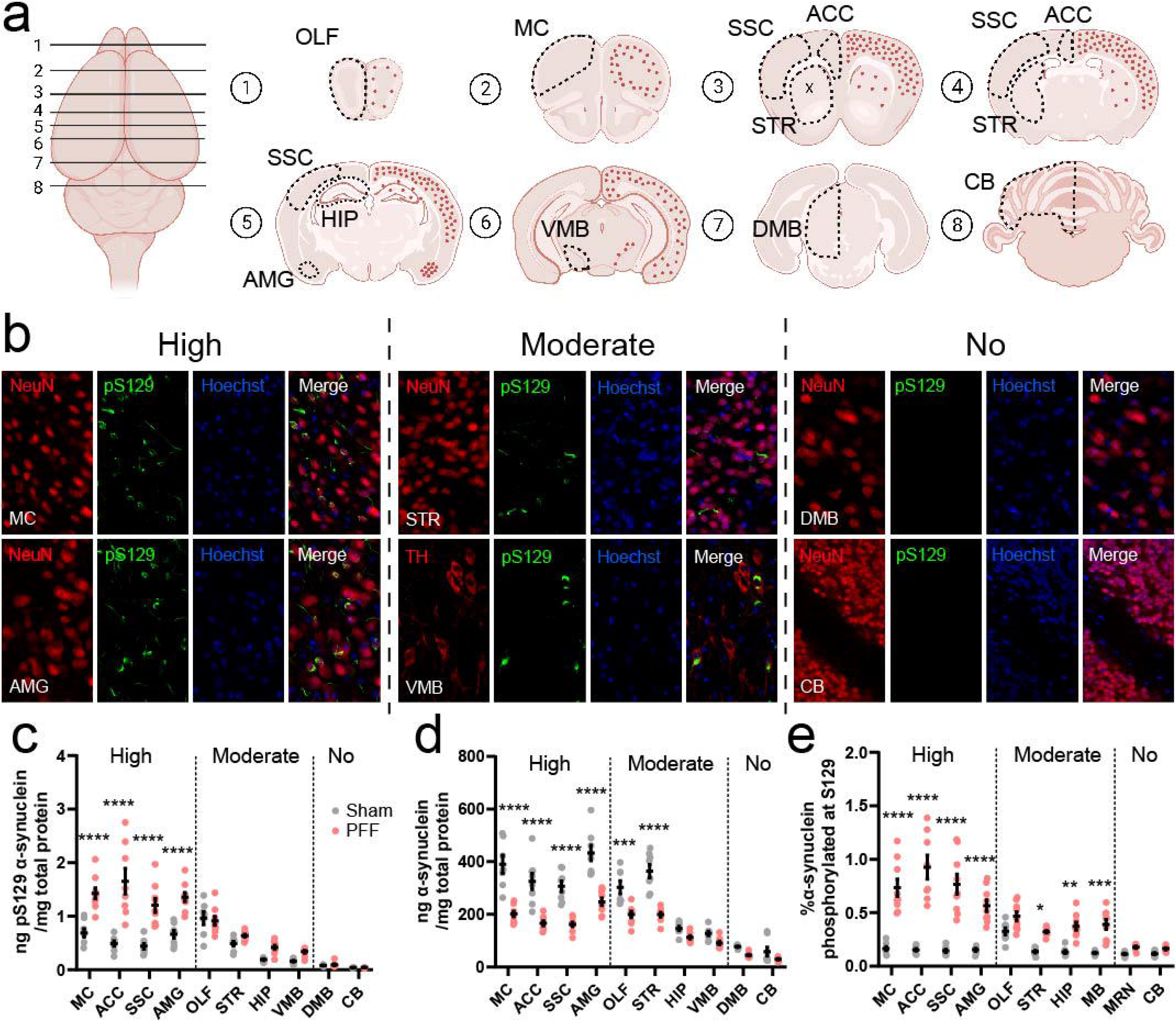
Profiling changes to the regional abundance of total and pS129 α-synuclein in the PFF mouse brain. The deposition of pS129 α-synuclein was highly heterogeneous throughout the brains of mice inoculated with mouse α-synuclein PFFs (**a, b**). Panel (**a**) depicts the representative burden of pS129 α-synuclein deposits (red dots) within the PFF mouse brain, which was assessed using pS129 α-synuclein immunofluorescent staining (EP1536Y, Abcam) of fixed 30µm free-floating coronal brain tissue sections from 3-month-old PFF mice (**b**). Full immunofluorescent characterization of all investigated brain regions in sham and PFF mice is presented in **Supplementary Figure 4**. Phosphorylated S129 (**c**) and total (**d**) α-synuclein was quantified in fresh tissue extracts from sham and PFF mouse brain regions exhibiting high (MC, ACC, SSC, AMG), moderate (OLF, STR, HIP, VMB) or no (MRN, CB) pS129 α-synuclein pathology using the reformulated SureFire Ultra pS129 and total α-synuclein assays. The proportion of pS129 α-synuclein (**e**) was calculated by normalising pS129 α-synuclein levels to the total amount of α-synuclein in these same tissue extracts. Data in panels **c-e** represent mean ± standard error of the mean. * *p*<0.05, ** *p*<0.01, *** *p*<0.001, **** *p*<0.0001; two-way ANOVA with Sidak’s multiple comparisons post-hoc tests. ACC, anterior cingulate cortex; AMG, amygdala; AU, arbitrary units; CB, cerebellum; DMB, dorso-medial midbrain; HIP, hippocampus; MC, motor cortex; OLF, olfactory bulb; SSC, somatosensory cortex; STR, striatum; VMB, ventral midbrain.

We observed substantial heterogeneity in absolute levels of pS129 and total α-synuclein across the sham mouse brain, with the highest levels of both species present in cortical and limbic regions that are susceptible to developing substantial pS129 α-synuclein pathology upon intrastriatal PFF inoculation (0.44-0.69ng pS129 α-synuclein/mg total protein, 306.7-433.8ng total α-synuclein/mg total protein). By contrast, phosphorylated and total α-synuclein levels were much lower in sham mouse regions that are spared from pS129 α-synuclein pathology following PFF inoculation, such as the DMB and CB (0.04-0.08ng pS129 α-synuclein/mg total protein, 58.1-76.7ng total α-synuclein/mg total protein) (**Figure 4c,d**). Despite this heterogeneity, the proportion of α-synuclein S129 phosphorylation was remarkably consistent (0.12-0.16%) across all investigated brain regions in sham mice (**Figure 4e**). The olfactory bulb constituted an exception to this observation, possessing a 2.4-fold higher proportion of α-synuclein S129 phosphorylation compared with all other sham mouse brain regions (0.33%; *p<*0.0001 vs all other brain regions, One-way ANOVA with Dunnett’s multiple comparisons post-hoc tests, *q =* 7.4-9.8, *DF* = 70), consistent with previous data^22^.

In whole tissue extracts from PFF-inoculated mice, absolute levels of pS129 α-synuclein were significantly higher in regions of high pathological burden (1.2-1.6 ng/mg total protein) compared with sham mice (**Figure 4c**), as was the proportion of α-synuclein S129 phosphorylation (0.57-0.93%) (**Figure 4e**). This was accompanied by a 43-49% reduction in total α-synuclein levels in these regions of PFF mice compared with sham mice (**Figure 4d**). Similar trends were identified in PFF mouse brain regions exhibiting moderate pathological burden, albeit to a lesser magnitude (0.3-0.9 ng pS129 α-synuclein/mg total protein; 0.32-0.47% pS129 α-synuclein; 23-45% reduction in total α-synuclein) (**Figure 4c-e**). No statistically significant change in any of these variables was identified in PFF mouse brain regions spared from pS129 pathology.

α-Synuclein is predominantly expressed within pre-synaptic nerve terminals^23,24^, hence the reduction in total α-synuclein in regions of high-moderate pS129 α-synuclein pathology may reflect reduced synaptic terminal densities in these regions. To address this possibility, we quantified levels of the pre-synaptic vesicular protein synaptophysin, which constitutes a robust proxy for pre-synaptic architecture and abundance that has been used previously to inform on synaptic density in α-synuclein PFF models^25,26^. Synaptophysin levels were unchanged in the OLF, ACC, AMG, STR and CB between PFF and sham mice (**Supplementary Figure 5**), suggesting reductions in total α-synuclein do not derive from reduced synaptic density in PFF mice.

### Altered subcellular distribution of total and pS129 α-synuclein in PFF mice

Changes to the compartmental distribution of pS129 α-synuclein have been previously reported in the PFF mouse brain^27,28^, however no study has been able to measure these changes due to the absence of quantitative α-synuclein assays. We addressed this knowledge gap by quantifying the distribution of total and pS129 α-synuclein in PBS-soluble (cytosolic, interstitial), triton-X(TrX-)soluble (membrane-bound), and SDS-soluble (aggregated) brain tissue fractions^29^ from PFF and sham mouse brain regions using our reformulated α-synuclein assays (**Supplementary Tables 9, 10**). The compartmental distribution of α-synuclein was highly heterogeneous between sham mouse brain regions (**Figure 5a-c**). While the majority (94.2-99.9%) was localized to the cytosol and interstitium (**Figure 5a**), membrane-bound α-synuclein was enriched 42-fold (5.7%) in regions prone to developing severe synucleinopathy, and 26-fold (3.4% total) in healthy brain regions that were susceptible to developing moderate synucleinopathy upon PFF inoculation (**Figure 5b**). Extremely little α-synuclein (<0.24%) was found to exist naturally as aggregated protein in sham mice (**Figure 5c**).

**Figure 5.**
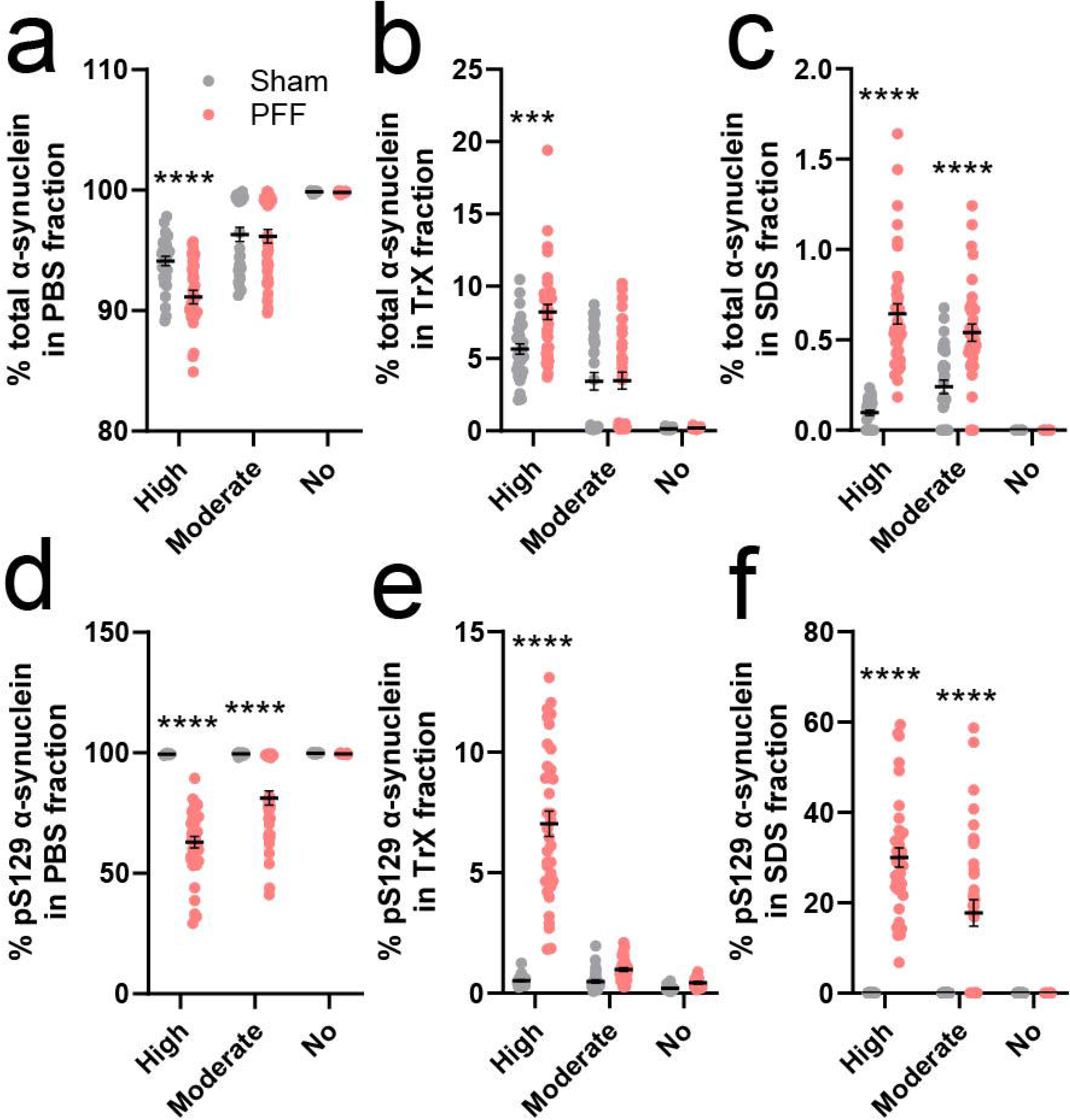
Characterising alterations to the compartmental distribution of total and pS129 α-synuclein in the PFF mouse brain. Fresh tissues from sham and PFF mouse brain regions exhibiting high (MC, ACC, SSC, AMG), moderate (OLF, STR, HIP, VMB) or no (MRN, CB) pS129 α-synuclein pathology following PFF inoculation were fractionated sequentially using saline solution containing protease and phosphatase inhibitors (PBS), followed by the same buffer containing 0.5% Tween-20 and 0.5% Triton X-100 (TrX), and finally with a TrX solution that had 2% SDS added to it (SDS). Total (**a-c**) and pS129 (**d-f**) α-synuclein were then quantified in these fractions using the new SureFire Ultra assays. Data in panels represent mean ± standard error of the mean. *** *p*<0.001, **** *p*<0.0001; two-way ANOVA with Sidak’s multiple comparisons post-hoc tests. Associated data are contained within **Supplementary Tables 9-10**. AU, arbitrary units.

Despite significant alterations to total α-synuclein abundance in some brain regions following PFF inoculation, the compartmental distribution of α-synuclein in PFF mice was only marginally different from sham mice in regions of severe pathological burden (**Figure 5a-c**). In these regions, intrastriatal PFF inoculation promoted a redistribution of 3.1% of cytosolic/interstitial α-synuclein (91.14% total) to cellular membranes (2.55% increase, 8.22% total) and protein aggregates (0.55% increase, 0.64% total). Total α-synuclein distribution in remaining brain regions was near identical between PFF and sham mice (**Figure 5a-c**), with 96.3-99.9% localized to the cytosol and interstitium, 0.1-3.4% bound to membranes and <0.3% contained within protein aggregates.

Contrast to total α-synuclein, the subcellular distribution of pS129 α-synuclein was remarkably similar between brain regions in sham mice (**Figure 5d-f**), being almost exclusively (99.5-99.8%) localized to the cytosol and interstitium (**Figure 5d**). This distribution became significantly disrupted upon PFF inoculation, with 17.8% of pS129 α-synuclein contained in the insoluble SDS-extractable fraction in brain regions exhibiting moderate synucleinopathy, which rose to 30.1% in regions exhibiting severe pS129 α-synuclein pathology (**Figure 5f**). Comparatively little (1.0-7.0%) pS129 α-synuclein was membrane-bound in these brain regions (**Figure 5e**).

### Aggregation of membrane-bound α-synuclein occurs largely in the absence of S129 phosphorylation

Aggregation of α-synuclein precedes S129 phosphorylation in PFF-inoculated mice^30^, although until now there have been no tools to quantify the magnitude, regional distribution, nor subcellular compartmentalization of these changes. We sought to develop new insights into the relationship between α-synuclein aggregation and S129 phosphorylation by profiling aggregated α-synuclein in the same tissue extracts as those used to measure total and pS129 α-synuclein. We first employed immunoblotting to assess alterations to the abundance of monomeric (**Figure 6a**) and SDS-resistant multimeric (dimers, trimers, oligomers) pS129 α-synuclein species (**Figure 6b**) in whole tissue extracts from PFF and sham mice. Brain regions exhibiting a high pathological burden in PFF mice (**Figure 4**) exhibited a significant reduction in monomeric pS129 α-synuclein compared with corresponding sham mouse regions, which was accompanied by a significant increase in multimeric pS129 α-synuclein. These changes were diminished in brain regions exhibiting moderate pS129 α-synuclein pathology and were absent in those lacking synucleinopathy. Pan-α-synuclein immunoblotting failed to detect multimeric α-synuclein in fractions where robust detection of pS129 α-synuclein multimers were present in the immunoblots (**Figure 6c,d**). The lack of multimeric α-synuclein species detectable by the total α-synuclein antibody suggests it only is a minor fraction of total α-synuclein that become phosphorylated on Ser129.

**Figure 6.**
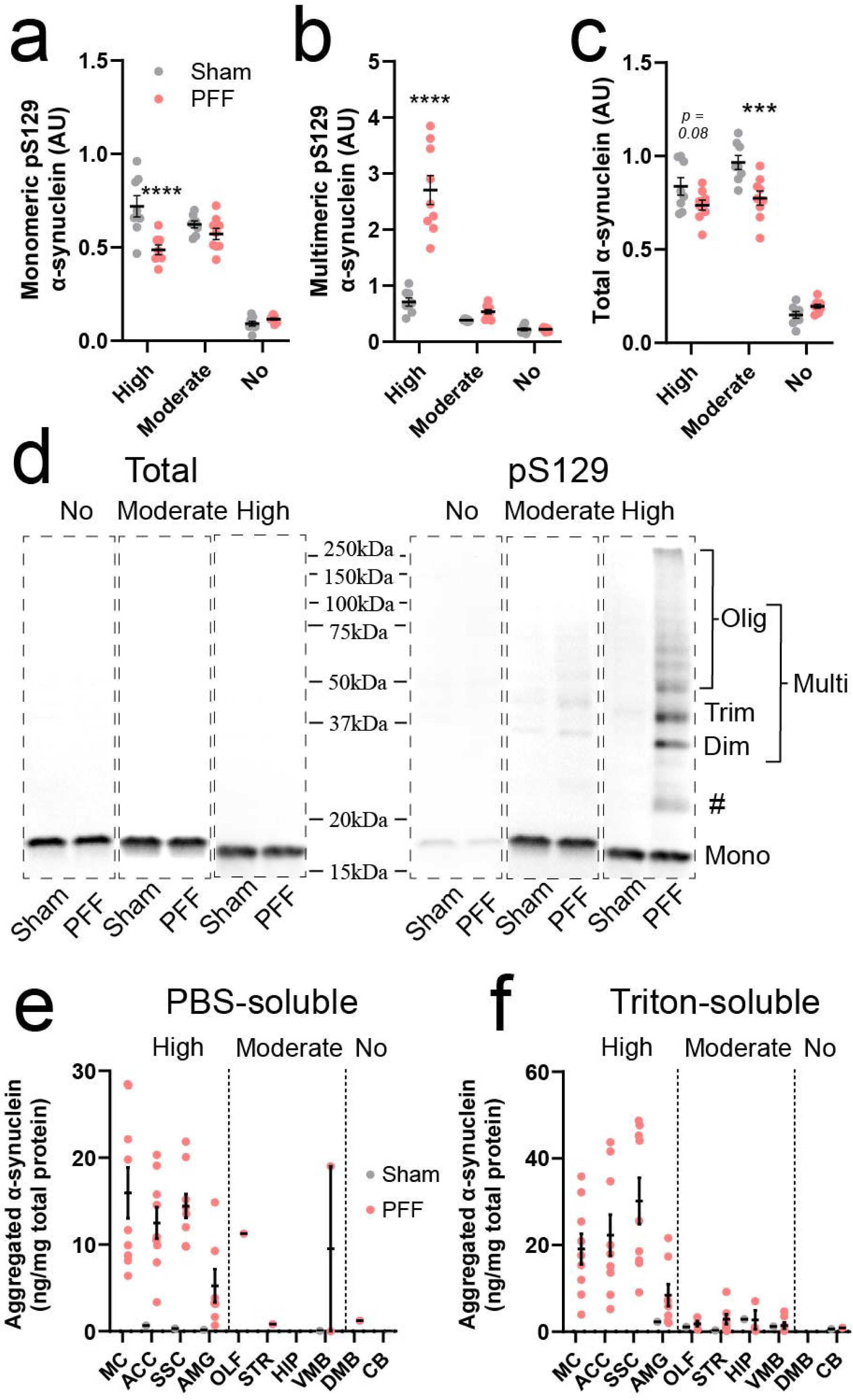
Profiling α-synuclein aggregation in the PFF mouse brain. Semi-quantitative analysis of monomeric (**a**) and multimeric (**b**) pS129 α-synuclein (D1R1R, Cell Signalling Technology^®^), as well as total α-synuclein (**c**; Syn1, BD Biosciences), was performed in whole tissue extracts from representative sham and PFF mouse brain regions exhibiting high (ACC), moderate (STR) or no (CB) pS129 α-synuclein pathology using immunoblotting (*n* = 8-9/group/category). Panel (**d**) depicts representative total and pS129 α-synuclein immunoblots from each representative brain region (ACC, STR, CB; full blots displayed in **Supplementary Figure 4**). All regions exhibited monomeric (Mono) native and pS129 α-synuclein in total and pS129 immunoblots, respectively. Phosphorylated S129 α-synuclein dimers (Dim), trimers (Trim) and Oligomers (Olig) were also observed in pS129 α-synuclein immunoblots, which we collectively termed multimeric (Multi) pS129 α-synuclein. Aggregated α-synuclein was quantified in PBS- (**e**) and triton-soluble (**f**) tissue fractions from all 10 brain regions of interest using the LEGEND MAX α-Synuclein Aggregate ELISA assay. Only datapoints above the assay’s lower limit of quantification (0.015ng/mL) were plotted, which for many groups meant plotting one or no values. Associated data are contained within **Supplementary Table 11**. Data in panels **a-c, e** and **f** represent mean ± standard error of the mean. # Phosphorylated S129 α-synuclein immunoreactivity just above the 20kDa molecular weight marker is consistent with glycosylated (O-GlcNAc) monomeric α-synuclein.*** *p*<0.001, **** *p*<0.0001; two-way ANOVA with Sidak’s multiple comparisons post-hoc tests. ACC, anterior cingulate cortex; AMG, amygdala; AU, arbitrary units; CB, cerebellum; DMB, dorso-medial midbrain; HIP, hippocampus; MC, motor cortex; OLF, olfactory bulb; SSC, somatosensory cortex; STR, striatum; VMB, ventral midbrain.

Next, we employed the LEGEND MAX α-Synuclein Aggregate ELISA (#448807, BioLegend^®^) to quantify the abundance of aggregated α-synuclein in PBS- (**Figure 6e**) and TrX-soluble (**Figure 6f**) tissue extracts from PFF and sham mice. No α-synuclein aggregation was detected above the assay’s lower limit of quantification (0.02 ng/mL) in 94% of samples from sham mice, with low (0.3-2.9 ng/mg total protein) aggregation detected in the remaining 10-of-160 fractions from all investigated brain regions. While data were similar for PFF mouse brain regions lacking pS129 α-synuclein pathology (0.9-1.2 ng/mg total protein), brain regions exhibiting a high pS129 α-synuclein burden contained up to 44.6 ng aggregated α-synuclein per mg of total protein, which was marginally more abundant in membrane-associated fractions (53.4-66.2% aggregates) compared with the cytosol/interstitium (33.8-46.6% aggregates). This differs significantly from regions of moderate pS129 α-synuclein pathology, where the majority of aggregated α-synuclein was membrane bound (78.3-100%), albeit at much lower quantities (2.7-4.6 ng/mg total protein).

## Discussion

Phospho-S129 α-synuclein is highly enriched in Lewy pathology^31^, hence technologies capable of high throughput measurement of pS129 α-synuclein are of great interest in synucleinopathy research. Several assays have been developed to measure human pS129 α-synuclein in human or transgenic mouse tissue extracts and biofluids^12,18,32-36^, however many of these cannot detect mouse pS129 α-synuclein or exhibit poor sensitivity for this target, presumably owing to discrepancies in amino acid residues at 7 sites across the protein^37^. This constitutes a critical unmet need given measurement of pS129 α-synuclein in non-transgenic mouse models is key to understanding the physiological function of α-synuclein and its aetiological contribution to synucleinopathies. By reformulating existing SureFire Ultra pS129 and total α-synuclein assays, we developed new ultrasensitive immunoassay technologies to quantify both mouse and human α-synuclein isoforms in mouse and human tissue and cell extracts. These assays are already commercially available (pS129, #ALSU-PASYN-B; total, #ALSU-TASYN-B; Revvity) and are designed as no-wash assays that can be completed within 2hrs of incubation time using as little as 1 µL brain tissue extract or 4 µL cell lysate (10x homogenization buffer volume (µL)/tissue weight (mg)) per triplicate pS129 and total α-synuclein measurement. Their calibrator matrix (assay buffer) is also highly suitable for measuring mouse and human pS129 and WT α-synuclein in mouse and human tissue and cell extracts, provided samples have been diluted beyond their MRDs. Exhibiting a significantly higher sensitivity (5-to-20-fold) and wider LDR (5-to-10-fold) compared with alternative human α-synuclein measurement technologies^12,18,19^, these assays are ideal for accurate and high-throughput sample processing using minimal material.

Application of reformulated SureFire Ultra assays to PFF and sham mouse brain tissues in this study clearly demonstrates their potential to parse valuable information from synucleinopathy disease models. The model of intrastriatal injection of α-synuclein PFF in wild type mice has become a mainstay in synucleinopathy research, recapitulating region-specific α-synuclein phosphorylation and aggregation observed in human disease^21^. It is unclear why select brain regions develop significant α-synuclein pathology above others and what role S129 phosphorylation plays in this process, largely as we have been unable to quantify mouse pS129 α-synuclein in these models in the past. While immunohistochemical characterization of pS129 synucleinopathy in PFF mice allows visualization and semi-quantitative estimation of relative pS129 α-synuclein burden across different brain regions, it cannot provide pS129 or total α-synuclein concentrations nor inform on the proportion of α-synuclein S129 phosphorylation across different brain regions. In this study, we demonstrate that these measurements are now made possible by reformulated SureFire Ultra α-synuclein assays, which match closely with qualitative immunohistochemical estimates of pS129 α-synuclein pathology across the 10 brain regions of interest. Regions of high, moderate and no pS129 α-synuclein immunofluorescence identified in fixed PFF mouse brain tissues can be clearly delineated using assay readouts in fresh brain tissue extracts (**Figure 4**), with sham and PFF mice also differentiated by the concentration and proportion of pS129 α-synuclein in regions of high pS129 α-synuclein burden. Analysing the same brain extracts with the BioLegend^®^ α-synuclein aggregate ELISA allowed us to demonstrate that the regions with the highest increases in pS129 α-synuclein burden also displayed large increases in both cytosolic and membrane-bound α-synuclein aggregates (Fig. 6e, f). This aggregation occurred alongside reductions in soluble pS129 α-synuclein, suggestive of intricate relationships between α-synuclein solubility, membrane binding, pS129 phosphorylation and conversion into SDS-insoluble aggregates. These data not only showcase the value of our pS129- and aggregate-assays in dissecting disease mechanisms in murine models, but also signify their potential in assessing the efficacy of treatments targeting α-synuclein phosphorylation or aggregation in preclinical murine models of synucleinopathy.

Aside from evaluating the performance of the new assays in mouse brain tissues, our data reveals novel biological insights into α-synuclein biology across the mouse brain and how this may influence the regional development of synucleinopathy upon PFF inoculation. Remarkable consistency in % α-synuclein S129 phosphorylation across most mouse brain regions in the WT mice highlights tight regulation of this modification under physiological conditions^38,39^. It is unclear whether its selective enrichment within the olfactory bulb signposts a differential regulation in this region. Others have suggested that the higher baseline pS129 α-synuclein point to the olfactory bulb as a regional nucleation point for the spread of synucleinopathy throughout the brain^22^, however our data indicate that this would have to occur in the absence of significant increases in total, proportional pS129 α-synuclein or aggregated α-synuclein in this region (**Figure 4c, e; Figure 6e, f**). Detailed examination of α-synuclein pathology across the PFF mouse brain at multiple ages post-inoculation is needed to comprehensively address this point. It must also be acknowledged that, while PFF mice are a suitable model for studying the development of synucleinopathy, they do not necessarily recapitulate the natural regional expression of this pathology across all human synucleinopathies alike, irrelevant of injection site.

A plausible interpretation of our data is that higher baseline total and pS129 α-synuclein concentrations in certain brain regions may underlie their susceptibility to developing moderate-to-severe synucleinopathy upon intrastriatal PFF inoculation. Indeed, increased concentrations of α-synuclein enhance the kinetics of fibrillization^40^, as a corollary, the development of synucleinopathy throughout the brain may be more dependent on regional α-synuclein concentrations rather than bio-distribution from the injection site. Further investigations into the development of synucleinopathy in mice following varied anatomical inoculations from the same PFF batch are warranted to assess the validity of this theory.

In addition to α-synuclein concentration, our data highlight a clear relationship between the protein’s subcellular compartmentalization and synucleinopathy load following PFF inoculation. Under physiological conditions, α-synuclein is thought to exist in a dynamic equilibrium between its natively unfolded cytosolic isoform and membrane bound species^41,42^. We determined that brain regions with naturally higher levels of membrane-bound α-synuclein are more susceptible to developing moderate-to-severe synucleinopathy following intrastriatal PFF inoculation. One possible explanation for this observation stems from data demonstrating that membrane-bound α-synuclein has a high aggregation propensity and can seed the aggregation of cytosolic α-synuclein^43^, although the trigger for this seeding remains unclear. Importantly, it is clear from our data that phosphorylation of S129 is not associated with α-synuclein membrane-binding under physiological conditions^44^, with only 0.2-0.5% of pS129 α-synuclein (<0.0017% total α-synuclein) bound to membranes across all investigated sham mouse brain regions. Furthermore, extremely little pS129 α-synuclein was membrane-bound in regions of moderate-to-severe synucleinopathy in PFF mice (<0.0085% total α-synuclein) despite a substantial increase in membrane-bound total α-synuclein in these regions.

## Conclusion

In summary, we reformulated existing SureFire Ultra pS129 and total α-synuclein assays to enable quantification of both mouse and human α-synuclein isoforms in mouse and human tissue and cell extracts with extremely high sensitivity. By combining these assays with a an aggregate α-synuclein ELISA, we highlight the ability of these assays to leverage novel biological insights into α-synuclein biology from established synucleinopathy mouse models by enabling quantification of both pS129 α-synuclein and aggregate pathology in these models. Application of these assays to PFF and sham mouse brain tissue fractions provided quantitative estimates of absolute, proportional pS129 α-synuclein and aggregate concentrations across the mouse brain under physiological and pathological conditions. These data support a model where α-synuclein aggregation in the PFF model is driven by higher baseline concentrations of membrane-bound α-synuclein in select brain regions, which may drive aggregation, as well as conversion to large insoluble aggregates and S129 phosphorylation. Further studies are needed to clarify the order of these molecular events. Future application of these new technologies to preclinical murine models of synucleinopathy employed in drug discovery studies also has the potential to improve outcome monitoring for therapies targeting pS129 and non-modified α-synuclein.

## Methods

### The AlphaLISA SureFire Ultra platform

For the purposes of this project, we obtained complete AlphaLISA SureFire Ultra kits containing all assay components as preformulated ready-to-use solutions. This not only minimized assay complexity, improved robustness and enhanced reproducibility, but more importantly will enable other investigators to obtain the same material from the supplier directly and perform their experiments as described herein. As detailed below, our work included assessment of multiple new formulations of SureFire Ultra Total α-synuclein and Phospho-α-synuclein (Ser129) kits, as well as previous formulations of these kits (PerkinElmer, Total α-synuclein; ALSU-TASYN-A, Phospho-α-synuclein (Ser129); ALSU-PASYN-A). What is described in this section applies to all tests that have been performed as well as the final product that has since been released by the manufacturer (Revvity, Total α-synuclein; ALSU-TASYN-B, Phospho-α-synuclein (Ser129); ALSU-PASYN-B).

The AlphaLISA SureFire Ultra is a high sensitivity, high throughput assay for the robust detection of analytes in biological samples. It is a no-wash immunoassay whose function is based on the luminescent proximity principal using oxygen-channelling chemistry, as shown in Supplementary Figure 1. In brief, its design utilizes two antibodies against a target protein, one of which is biotinylated and the other of which is conjugated to a proprietary CaptSure^TM^ tag. The differential tagging of these antibodies enables them to selectively bind to one of two types of Alpha beads; Alpha donor beads are streptavidin-coated to allow capture of the biotinylated antibody, whilst Alpha acceptor beads are coated with a proprietary CaptSure^TM^ agent to allow binding of the respectively tagged antibody. Once all components have been added to the sample and both antibodies and beads are complexed with the target protein, the sample is illuminated using 680nm light, causing the release of singlet oxygen molecules from the donor beads, triggering the acceptor beads in close proximity (<200 nm) to emit a signal at 615 nm. This emission intensity is directly proportional to the amount of the target protein within the assay’s dynamic range.

The assay was performed according to the manufacturer’s recommended protocol, with the exception that the positive control lysate supplied with each kit was replaced by purified human or mouse WT α-synuclein, or pS129 α-synuclein protein standards, which are described below. Assay signals were measured using an EnVision^TM^ multimodal plate reader under the manufacturer’s preset AlphaScreen^TM^ protocol.

### Recombinant α-synuclein protein standards

Mouse (#RP-009) and human (#RP-003) WT α-synuclein, as well as human pS129 α-synuclein (#RP-004), were obtained from Proteos (Kalamazoo, MI, USA), where they were produced in collaboration with the Michael J Fox Foundation for Parkinson’s Research. Recombinant mouse and human WT α-synuclein protein standards were provided as 100 µL stock aliquots of 10 mg/mL α-synuclein in 10 mM Tris, 50 mM NaCl, pH 8, which were diluted to 1 mg/mL intermediate working aliquots using 900 µL of 10mM Tris, 50mM NaCl, pH 8 containing 0.55% Tween-20, 0.55% Triton X-100, 0.55% BSA and 0.055% sodium azide (final concentrations: 10 mM Tris, 50 mM NaCl, pH 8 containing 0.5% Tween-20, Triton X-100 and BSA, 0.05% sodium azide). Recombinant human pS129 α-synuclein was provided as 1 mg aliquots of lyophilized standard, which were reconstituted to 1 mg/mL intermediate working aliquots using 1 mL of 10 mM Tris, 50 mM NaCl, pH 8 containing 0.5% Tween-20, Triton X-100 and BSA, 0.05% sodium azide.

Mouse pS129 α-synuclein was not available from the same supplier. We therefore produced it in-house from a 1.25 mg/mL solution of mouse WT α-synuclein in kinase reaction buffer, which was generated by diluting a 100 µL (10 mg/mL) aliquot of this standard 8-fold in 700 µL of 22.86 mM HEPES, 1.25 mM ATP, 2.29 mM DTT, 11.43 mM MgCl2, pH 7.4 (final concentrations: 20 mM HEPES, 1.09 mM ATP, 2 mM DTT, 10 mM MgCl2, pH 7.4). Phosphorylation was induced by incubating 1 µg of recombinant human polo-like kinase 3 (PLK3) protein (Thermo Fisher Scientific, MA, USA; #PR7316B) with 50 µL of mouse WT α-synuclein in kinase reaction buffer for 8 hrs at 30°C. Immunoblotting and electrospray ionisation mass spectrometry were used to confirm complete phosphorylation of α-synuclein S129 (**Supplementary Figure 3)**. All standards were stored at -80°C in 10 µL aliquots, with the exception of lyophilized human pS129 α-synuclein, which was stored at -20°C as per manufacturer’s recommendations. Any unused standard leftover from a thawed aliquot was discarded at the end of each assay working day.

### HEK293 cell culture and transfection

Wild-type (#ab255449) and α-synuclein KO (#ab255433) HEK293 cells were obtained from Abcam, while PLK3-HEK293 cells were produced by transfection of wild-type HEK293 cells with Myc-DDK-tagged human polo-like kinase 3 (PLK3; # RC203352, OriGene, MA, USA). All experimental procedures conducted on these cell lines were approved by the University of Sydney Institutional Biosafety Committee (#21E012).

For transfection experiments, plasmids were first transformed into One Shot TOP10 Chemically Competent *Escherichia coli* (Thermo Fisher Scientific; #C404003) according to manufacturer’s instructions. Selection for successful transformation was performed using Ampicillin (100 µg/mL, control) or Kanamycin (25 µg/mL, PLK3) prior to the expansion of resistant colonies in 100 mL of Luria-Bertani (LB) broth and purification of plasmid DNA using a PureLink HiPure Plasmid Filter Midiprep Kit (Thermo Fisher Scientific; #K210014), according to manufacturer’s instructions. Aliquots of transformed bacteria were stored in a 1:1 mixture with 50% glycerol at -80°C, while purified plasmid DNA was stored reconstituted in deionized water at -20°C. Wild-type and α-synuclein KO HEK293 cells were then seeded into 6-well plates (45 x 10^4^ cells/well) before being transfected with 5 µg DNA per well using Lipofectamine 3000 (Thermo Fisher Scientific; #L3000015) according to manufacturer’s instructions. Cells in were grown to 90% confluency before being harvested, washed in 1xPBS, pelleted, and stored until use at -80°C.

For assay validation experiments, transfected and non-transfected cells were grown to 90% confluency in 6-well plates using a 1:1 mixture of Dulbecco’s Modified Eagle Medium and Nutrient Mixture F-12 (DMEM/F-12, Thermo Fisher Scientific; #11320033) supplemented with 10% fetal bovine serum (heat inactivated, Thermo Fisher Scientific; #16000044), 1% GlutaMAX (Thermo Fisher Scientific; #35050061) and 1% penicillin streptomycin (10,000 U/mL, Thermo Fisher Scientific; #15140122), before being harvested, washed in 1xPBS and pelleted. Cell pellets were resuspended in 1x assay lysis buffer containing protease (Sigma; #11697498001) and phosphatase (Sigma; #4906845001) inhibitors (10 volumes (µL) per mg cell pellet) using a Branson SFX250 Sonifier (Emerson, St. Louis, MO, USA; #101-063-965R) equipped with a 3/32” probe microtip. Pellets were sonicated on ice for 2 minutes with a 5 sec pulse (70% amplitude), followed by a 25 second pause.

### Wild-type and α-synuclein knock-out mice

Female WT C57BL/6 mice (12 weeks old) were obtained from the Animal Resources Centre (Canning Vale, WA, Australia) and housed within the Laboratory Animal Services facility at the Charles Perkins Centre (University of Sydney, NSW, Australia), with ad libitum access to food and water on a 12 h light-dark cycle at temperatures between 20-24°C and humidity between 40-70%. All experimental procedures were approved by the University of Sydney Animal Ethics Committee (#2021/2015).

Brain tissues were excised from wild-type and α-synuclein KO mice following intraperitoneal injection with a lethal dose of sodium pentobarbitone (Lethabarb, 100mg/kg) and transcardial perfusion with ice-cold 0.9% NaCl for 10 mins at a constant flow rate of 6mL/min using a peristaltic pump (World Precision Instruments, Sarasota, FL, USA). Brains were then sagittally bisected, with one hemisphere frozen immediately on dry ice and stored at -80°C. The contralateral hemisphere was cut into four pieces of approximately equal size, which were then sonicated for 3 mins or until no visible tissue pieces remained using the same microtip-based sonication protocol described above for cell extracts. These four homogenates were then combined to create a whole brain homogenate, which was centrifuged at 20,000 *g* for 30mins at 4°C before the supernatant was collected and aliquoted for subsequent analyses.

Homozygous *Snca* knock-out mice were obtained from Jackson Laboratories (SNCA^-/-^; C57BL/6N-Snca^tm1Mjff^/J; JAX stock #016123)^45^. They were bred under the breeding permit 2022-15-0202-001135 issued by the Danish Veterinary and Food Administration. Animals were housed with light-dark cycles of 12 h intervals. Mice were fed standard chow and water ad libitum. Brain tissues were excised from 12-week-old wild-type and α-synuclein KO mice following intraperitoneal injection with a lethal dose of sodium pentobarbitone (Lethabarb, 100 mg/kg) and transcardial perfusion with ice-cold 0.9% NaCl for 10mins at a constant flow rate of 6mL/min using a peristaltic pump (World Precision Instruments, Sarasota, FL, USA). Upon removal, brains were dissected sagittally down the midline and snap-frozen in liquid nitrogen.

### SureFire assay validation

Five new total and pS129 assay antibody pairings were screened against purified mouse and human WT and pS129 α-synuclein protein standards, which we named as total or pS129 α-synuclein assay pair 1-5 in this publication. Antibody pairs were all designed to contain at least one antibody known to robustly detect mouse and human α-synuclein in fixed and/or frozen tissue sections and protein extracts using immunohistochemistry and/or immunoblotting. Standards were prepared for assay screening by serially diluting 1 mg/mL intermediate working aliquots of each standard to concentrations between 2.24 ng/mL and 0.038 pg/mL using 1x Alpha SureFire Ultra Lysis Buffer (Revvity, MA, USA; #ALSU-LB-100ML), hereon referred to simply as assay lysis buffer. Serial dilutions for a given standard were all performed using the same pipette tip and were conducted in Protein LoBind^®^ Tubes (Eppendorf, Hamburg, Germany; #0030108434) to minimize loss of α-synuclein between dilutions through adherence to pipette tips and tube walls. Each dilution of every standard was measured in 3 technical replicates and data expressed as mean ± standard deviation to convey average assay signal and intra-assay variation. We defined the limit of detection (LoD) and lower limit of quantification (LLoQ) as 3 and 6 standard deviations above the mean of the blank, respectively. Inter-assay variability for each dilution of every standard was determined across three individual standard curves measured on separate days and was calculated by dividing the standard deviation of the replicates by their mean, multiplied by 100 to produce percentage variability.

The 2 formulations of each assay with the highest sensitivity for mouse and human isoforms of their target protein were then applied to serially diluted WT and α-synuclein KO mouse brain tissue supernatant and HEK293 cell lysates, as well as PLK3-HEK293 cell lysates. Each dilution of every sample was measured in 3 technical replicates and assay data (intra- and inter-assay variability, LoD, LLoQ) calculated as described for purified standards above. All subsequent experiments only employed the total and pS129 α-synuclein assay formulation exhibiting the highest sensitivity.

For parallelism experiments, WT and α-synuclein KO mouse brain tissue extracts were each pre-diluted 10- and 100-fold in assay lysis buffer, representing dilution factors above and below the MRD for mouse brain matrix. These pre-diluted WT extracts were then serially diluted up to 4096-fold using either assay lysis buffer or the equivalent dilution of α-synuclein KO extract, i.e. 10-fold diluted WT extract was diluted up to 4096-fold using 10-fold diluted KO extract. Wild-type and α-synuclein KO HEK293 cell lysates were processed in a similar manner, being pre-diluted 5- and 20-fold in assay lysis buffer.

Spike-in experiments were designed to assess recovery of WT and pS129 mouse α-synuclein from mouse brain tissue matrix. Wild-type mouse brain tissue extract was diluted 2000-fold in assay lysis buffer to ensure signal from endogenous α-synuclein was within the assay’s LDR, whereas α-synuclein KO extract was diluted 100-fold in assay lysis buffer to ensure minimal signal distortion by non-specific binding to matrix components (MRD – 24-fold). Wild-type and pS129 mouse α-synuclein standards were then spiked separately into diluted WT and KO mouse brain extracts to final estimated concentrations ranging between 10-640pg/mL (WT protein) and 0.625-40pg/mL (pS129 protein). These concentration ranges were designed to fall within the LDR of the total and pS129 α-synuclein assay, respectively. Wild-type mouse α-synuclein spike recovery was assessed with the total assay and pS129 mouse α-synuclein spike recovery was assessed with the pS129 assay. Spike concentrations were then measured in triplicate. Percentage spike recovery was calculated by dividing the mean measured spike concentration by the intended spike concentration and multiplying by 100. Measured spike concentrations for WT mouse α-synuclein were determined using the total α-synuclein kit standard curve for WT mouse α-synuclein, while those for pS129 mouse α-synuclein were determined using the pS129 α-synuclein kit standard curve for pS129 mouse α-synuclein.

### Mouse α-synuclein pre-formed fibril (PFF) generation and characterization

Wild-type monomeric mouse α-synuclein was produced as previously described^46^. Lyophilized wild-type monomeric α-synuclein was reconstituted in PBS pH 7.4 (Gibco, Billings, Montana, USA) and passed through a 100kDa Amicon^®^ Ultra Centrifugal Filter (Merck, Rahway, NJ, USA) to separate unwarranted oligomeric species, before being sterile filtered through a 0.22 µm filter (Merck) to remove additional particulates and microorganisms. Protein concentration was determined using a Pierce^TM^ BCA Protein Assay Kit (Thermo Fisher Scientific^TM^) according to manufacturer’s instructions and adjusted to 1.025 mg/mL with PBS. Next, 25 µL of 2 mg/mL sonicated α-synuclein pre-formed fibrils (produced previously^47^) was added to 975 µL of the 1.025 mg/mL monomeric α-synuclein solution to seed aggregation, at a concentration of 5% fibrils by mass%. The samples were incubated at 37°C for 72 h on an orbital shaker (1050 rpm). Samples were centrifuged at 15,600 g for 30min to pellet insoluble fibrils, which were subsequently resuspended in PBS. The protein concentration in this suspension was again determined using a Pierce^TM^ BCA Protein Assay (Thermo Fisher Scientific) and adjusted to 2 mg/mL using PBS. Fibrils were then sonicated for 20 min with 30 ms pulses at 30% power, followed by 70 ms pauses, using a Branson SFX250 Sonifier equipped with a 1” cup horn (Branson; #101-147-046), before aliquots were stored at -80°C. Amyloid structure of fibrils was confirmed using a Thioflavin-T binding assay as previously described (**Supplementary Figure 6**)^48^, while fibril fragment size (24nm) was measured by dynamic light scattering using a DynaPro^TM^ NanoStar^TM^ (Wyatt Technology, Goleta, CA, USA) and was not found to change following freeze/thaw (**Supplementary Figure 6**). Fibrils were also subjected to sedimentation analysis, whereby sonicated PFFs and monomeric standards were centrifuged at 25,000 g for 30 mins. Both the pellet and supernatant were then collected, the pellet resuspended in PBS, and immunoblot loading buffer added to both fractions. Samples were then incubated at 95°C for 10mins, before being loaded into pre-cast gels and subjected to electrophoresis as described above. Proteins were stained within gels using Coomassie Blue and imaged using a Fuj LAS-3000 Intelligent Dark Box (Fujifilm, Japan). Approximately 50% of the PFFs became soluble after sonication and freeze/thaw (**Supplementary Figure 6**), which we attributed to small fragment size given dynamic light scattering data did not indicate monomerization following freeze/thaw.

### Wild-type mouse stereotaxic injections

C57BL/6J mice weighing approximately 20 g were given prophylactic Temgesic (0.03 mg/kg) intraperitoneally fifteen minutes prior to anaesthesia induction with inhalable isoflurane (2-5%). Once anaesthesia was achieved, the dorsal head surface was shaved and animals secured to the stereotaxic frame (Stoelting, Wood Dle, IL, USA), fitted with a mouse face mask adapter. A 5 mm midline incision was made on the dorsal surface, the tissue retracted and bregma located. Animals received bilateral 2.5 µL intracerebral injections through a drilled burrhole of either 2 mg/mL mouse α-synuclein PFF’s (PFF; n=12) or sterile PBS (pH 7.4, sham; n=9) targeting the centre of the striatum (relative to bregma: +0.2 mm anteroposterior, ± 2 mm mediolateral, -3.2 mm dorsoventral from the dural surface). Injections were performed with a glass cannula (50 μm internal diameter) attached to a 10 μL Hamilton syringe (Hamilton Company, NV, USA) and injected at a constant dose rate of 0.4 μL/minute using an automated syringe pump (UltraMicroPump3, World Precision Instruments, FL, USA). The cannula was kept in-situ for 5 minutes at each target site prior to retraction and flushing with 0.9% sterile saline between injections. Following surgery, skin incisions were closed with suture and animals monitored closely during recovery.

### PFF mouse brain tissue harvesting and processing

Sham and PFF mice (*n* = 3/group) culled for histological profiling of pS129 α-synuclein burden 3 months post-intrastriatal PFF inoculation were injected intraperitoneally with a lethal dose of sodium pentobarbitone (Lethabarb^®^, 200 mg/kg), before being transcardially perfused with 0.9% NaCl (8mins, 6mL/min) followed by ice-cold 4% paraformaldehyde (PFA in 0.1 M phosphate buffer, pH 7.4, 8 mins, 6 mL/min) using a peristaltic pump (World Precision Instruments). Brains were excised whole, post-fixed in 4% PFA overnight at 4°C and transferred into 30% sucrose (w/v in 1xPBS) for 48 hrs at 4°C. Brains were then flash frozen using isopentane cooled to between -50°C and -60°C on dry ice, before being mounted onto microtome chucks using optimal cutting temperature compound. Whole mounted brains were sectioned coronally into 12x 30 µm section series’ and stored in antifreeze solution (30% glycerol, 30% ethylene glycol in 1x phosphate buffer (PB; pH 7.4)) at -20°C.

Sham and PFF mice (*n* = 8-9/group) culled for biochemical analyses 3.5 months post-intrastriatal PFF inoculation were lethally anaesthetized and transcardially perfused with 0.9% NaCl as above, before brains were harvested whole and bisected along the mid-sagittal axis. The right hemisphere was post-fixed in 4% PFA overnight at 4°C, transferred into 70% ethanol overnight at 4°C, and embedded in paraffin wax. The left hemisphere was dissected fresh to obtain tissue from 10 brain regions – the olfactory bulb (OLF), motor cortex (MC), anterior cingulate cortex (ACC), somatosensory cortex (SSC), amygdala (AMG), striatum (STR), hippocampus (HIP), ventral midbrain (VMB), cerebellum (CB) and dorso-medial midbrain (DMB). These specific brain regions were chosen for further biochemical examination as they exhibited high (MC, ACC, SSC, AMG), mild-moderate (OLF, STR, MB, HIP) or no (CB, DMB) pS129 α-synuclein burden in histological analyses of PFF brains at 3 months post-PFF inoculation.

Fresh tissues were suspended in 1x assay lysis buffer containing protease (Sigma; #11697498001) and phosphatase (Sigma; #4906845001) inhibitors (10 volumes (µL) per mg tissue) using a Branson SFX250 Sonifier (Emerson, St. Louis, MO, USA; #101-063-965R) equipped with a 3/32” probe microtip. Pellets were sonicated on ice for 2 minutes with a 5 sec pulse (70% amplitude), followed by a 25 second pause. One quarter of the total homogenate volume was then aliquoted and stored at -80°C (“whole tissue extract”), while the remainder was centrifuged at 20,000 *g* for 30mins at 4°C and the supernatant collected and designated the “PBS-soluble fraction”. PBS-insoluble pellets were then resuspended in 10 volumes (µL) of PBS fraction buffer supplemented with 0.5% Tween-20 and 0.5% Triton X-100 (TrX) using probe tip sonication (2x 5sec pulses, 70% amplitude, 25sec intermittent pause) and incubated on ice for 1 hr. These solutions were then centrifuged at 20,000 *g* for 30mins at 4°C and the supernatant collected and designated the “TrX-soluble fraction”. Any remaining TrX-insoluble pellets were fully dissolved in TrX fraction buffer supplemented with 2% SDS using probe tip sonication as described for the TrX fraction, and were designated the “SDS-soluble fraction”. Protein concentrations in PBS, TrX and SDS fractions were determined using a Pierce^TM^ BCA Protein Assay (Thermo Fisher Scientific).

### Immunofluorescence

Fixed tissue sections from sham and PFF mice were prepared for fluorescent staining and microscopy by first washing in PBS containing 0.1% Tween-20 (PBS-T) to remove antifreeze solution, before being subjected to antigen retrieval in citrate buffer (10mM sodium citrate, 0.05% Tween-20, pH 6.0) at 70°C for 30 mins. Sections were then washed again with PBS-T before being blocked (4% BSA (w/v), 1% casein (w/v), 1.5% glycine (w/v), 0.25% Triton X-100 (v/v) in 1x PBS) for 1hr at room temperature and incubated overnight at 4°C with primary antibodies directed against pS129 α-synuclein (rabbit, 1:10,000, EP1536Y, Abcam; #ab51253), tyrosine hydroxylase (chicken, 1:5,000, Abcam; #ab76442) and NeuN (rat, EPR12763, Abcam; #ab279297) diluted in blocking serum. Primary antibodies were detected using goat anti-host IgG secondary antibodies conjugated to AlexaFluor^TM^ 405, 488 or 647 dyes (1:1000 diluted in blocking solution), before sections were mounted onto microscope slides coated with gelatin-chrom alum and coverslipped using #1.5 coverslips and ProLong^TM^ Diamond Antifade Mountant (Thermo Fisher Scientific). Images were collected using a C2 laser scanning confocal microscope equipped with 405, 488 and 640 laser lines (Nikon, Minato-ku, Tokyo, Japan), as well as a Nikon Plan Apo VC 20x (0.75 DIC N2, 0.17 WD 1.0) and a Nikon CFI Plan Apochromat Lambda D 40x objective (0.95 DIC N2, 0.21 WD 1.0), Images were viewed and analysed using Fiji software (National Institute of Health, Bethesda, Maryland, USA).

### SureFire assay measurements in PFF mouse brain extracts

We first identified optimal dilution factors required to bring total and pS129 α-synuclein levels in each synucleinopathy mouse model, extract, and/or fraction, to within the LDRs of the total and pS129 assay (**Supplementary Table 8**). Whole tissue extracts (homogenates), as well as PBS-, TrX- and SDS-soluble brain tissue fractions, were diluted in assay lysis buffer to their optimal dilution factors where total and pS129 α-synuclein were measured. All dilution factors were beyond the MRD (24-fold) required to alleviate mouse brain matrix effects. Total α-synuclein concentration in diluted samples was calculated using the total α-synuclein kit standard curve for WT mouse α-synuclein. Phosphorylated S129 mouse α-synuclein was determined using the pS129 α-synuclein kit standard curve for pS129 mouse α-synuclein. Concentrations in diluted samples were then multiplied by their dilution factors and expressed as ng/mg total protein to account for any differences in protein concentration in original extracts. The proportion of α-synuclein phosphorylated at S129 was calculated by dividing ng pS129 α-synuclein/mg total protein by the ng total α-synuclein/mg total protein, expressed as a percentage.

### Immunoblotting

Immunoblotting for total and pS129 α-synuclein in protein standards (500ng) or tissue/cell extracts (20µg total protein) was performed by incubating samples in loading buffer (100 mM dithiothreitol (DTT), 3% SDS, 10% glycerol, 0.05% bromophenol blue, 62.5 mM Tris-base, pH7.4; Sigma-Aldrich, MO, USA) at 95°C for 10 mins before being loaded into 4-20% Mini-PROTEAN^®^ TGX^TM^ Precast Protein Gels (15 wells, 15uL; Bio-Rad, CA, USA) and subject to electrophoresis on a Bio-Rad Mini-PROTEAN^®^ Tetra Cell system (180 V, 40 mins, 4°C). Proteins were then transferred to Immun-Blot^®^ PVDF membrane (0.2 µm pore size; Bio-Rad) at 30 V for 2 hrs at 4°C, before membranes were fixed in 4% paraformaldehyde for 30mins at room temperature, washed in 1xPBS containing 0.1% Tween (PBS-T) and blocked in 5% bovine serum albumin (BSA, Sigma-Aldrich, #A7030) diluted in PBS-T for 1 hr at room temperature. Membranes were incubated overnight at 4°C with a mouse pan α-synuclein primary antibody (Syn1, BD Biosciences, NJ, USA; #610786) diluted 1:5,000 in blocking solution, before being washed with PBS-T and incubated for 2 hrs at room temperature with 1:5,000 goat anti-mouse horseradish peroxidase-conjugated secondary antibody (Thermo Fisher Scientific; #G21040) diluted in blocking solution. Protein signals were developed using ECL western blotting substrate (Bio-Rad) and detected using a Chemi-Doc^TM^ XRS imaging system (Bio-Rad) according to manufacturer’s instructions. Membranes were then incubated in stripping buffer (25mM glycine, 2% SDS, pH 2.0) for 30mins at room temperature, before being washed, blocked, probed and imaged as above using a rabbit pS129 α-synuclein-specific primary antibody (D1R1R, 1:5,000, Cell Signalling, MA, USA; #23706) and a goat anti-rabbit horseradish peroxidase-conjugated secondary antibody (1:5,000, Thermo Fisher Scientific; #G21234).

For quantitation of synaptophysin in mouse brain extracts, 7µg of total protein was incubated in loading buffer as described above and loaded into AnykD Mini-PROTEAN TGX Precast Protein Gels (15 wells, 15uL; Bio-Rad). Electrophoresis and protein transfer were performed as above, without membrane fixation, before membranes were dried overnight 2 h at room temperature and proteins stained with Sypro Ruby Protein Blot Stain (ThermoFisher Scientific), according to the manufacturer’s instructions. Sypro Ruby-stained membranes were imaged using a Chemi-Doc^TM^ XRS imaging system (630BP30nm filter, Bio-Rad), and total protein in each sample quantified by densitometry using ImageLab software (v.5.2, Bio-Rad). Following total protein quantification, membranes were blocked in 5% skim milk diluted in PBS-T for 1 hr at room temperature and incubated overnight at 4°C with a rabbit synaptophysin primary antibody (YE269, Abcam, Cambridge, UK; #ab32127) diluted 1:2,000 in blocking solution. Membranes were then washed with PBS-T and incubated for 2 hrs at room temperature with 1:3,000 goat anti-rabbit horseradish peroxidase-conjugated secondary antibody (Thermo Fisher Scientific; #G21234) diluted in blocking solution. Protein signals were developed and detected as above, and synaptophysin protein levels normalized to total protein levels within each sample to correct for variations in protein loading, as well as an internal standard (pooled from all tissue samples) to correct for variability between gels and immunoblot runs.

### α-Synuclein aggregate quantification

Aggregated α-synuclein was quantified in PBS- and TrX-soluble tissue fractions (5µg total protein/well) using the LEGEND MAX^TM^ α-Synuclein Aggregate ELISA (BioLegend) according to manufacturer’s instructions.

### Statistical analyses

Statistical analyses were performed using IBM SPSS Statistics (Version 27, IBM, Armonk, New York, United States) and GraphPad Prism (Version 7.02, GraphPad, San Diego, CA, USA). Parametric tests or descriptive statistics with parametric assumptions (standard two-way and one-way ANOVA, Pearson’s r) were used for variables meeting the associated assumptions, with data normality assessed using either the D’Agostino-Pearson (omnibus K2) normality test or the Shapiro-Wilk (Royston) normality test. Two-way ANOVAs were paired with Sidak’s multiple comparisons post-hoc tests to assess pair-wise comparisons between select diagnostic groups for a given variable. No outliers were detected using the combined robust regression and outlier removal (ROUT) method with a maximum false discovery rate of 5%. A *p*-value of less than 0.05 was accepted as the level of significance. Details of the statistical tests employed (test statistics, sample sizes, *p* values) for each variable of interest are included in the corresponding figure legend, results text or supplementary data section. An *n* of 8-9 cases per diagnostic group was strongly powered to detect differences in %pS129 α-synuclein (90-100% power, two-tailed *t-*test, α = 5%; SPSS software, IBM, Armonk, NY, USA) based on a preliminary power analysis of pilot data describing this measure in PFF- vs sham-inoculated WT mice (*n* = 3/group).

## Data Availability

All assay validation data are available in the main text or Supplementary Materials of this article. Any additional datasets generated using our experimental mouse models are available from the corresponding author on reasonable request.

## Acknowledgements

This study was funded by the Research Tools Program of the Michael J Fox Foundation for Parkinson’s Research (MJFF-021836), P.H.J was supported by Lundbeck Foundation grants R223-2015-4222, R248-2016-2518 for Danish Research Institute of Translational Neuroscience-DANDRITE, Nordic-EMBL Partnership for Molecular Medicine. For the purpose of open access, the authors have applied a CC BY public copyright license to all Author Accepted Manuscripts arising from this submission. The authors would like to acknowledge Assoc. Prof. Thomas J. D. Jørgensen (Department of Biochemistry and Molecular Biology, University of Southern Denmark, Denmark) for his insights into α-synuclein phosphorylation, as well as Yu Kwan (Charles Perkins Centre, The University of Sydney, Australia) and Benedicte Vestergaard (Department of Biomedicine, Aarhus University, Denmark) for their technical support. They also thank the Laboratory Animal Services Facility at the University of Sydney for their technical support in housing and maintaining animals used in this study.

## Author contributions

D.K., P.H.J. and N.D. conceived the study. D.K., P.H.J. and A.P. acquired ethical approvals. B.G.T., C.J.W., A.R., L.C., N.M.J. and H.G. collected primary data. B.G.T., D.K. and P.H.J. analyzed primary data and drafted the manuscript. All authors edited the manuscript.

## Additional information

**Supplementary information** is contained within the online version of this manuscript.

**Correspondence** and requests for data or materials should be addressed to Deniz Kirik or Poul Henning Jensen.

