## Supplementary Data for "Novel tools to quantify total, phospho-Ser129 and aggregated alpha-synuclein in the mouse brain"

### SUPPLEMENTARY INFORMATION

#### *Inventory of Supplementary Information*

Supplementary Figures 1-8

1) AlphaLISA SureFire Ultra assay platform and workflow schematics.
2) Characterization of the linear dynamic range of new SureFire Ultra assays using purified protein standards.
3) Evaluation of matrix effects in HEK293 cell lysate using a parallelism experiment.
4) Immunohistochemical profiles of pS129 α-synuclein across brain regions of interest in sham and PFF mice.
5) Immunoblot quantification of synaptophysin in sham and PFF mouse brain regions.
6) Full representative immunoblots characterising α-synuclein phosphorylation in brain tissue extracts from sham and PFF mice.
7) Full representative immunoblots characterising S129 phosphorylation in purified protein standards.
8) Characterisation of mouse pre-formed fibril (PFF) structure and solubility.

Supplementary Tables 1-11

1) Characterization of previous total and pS129 α-synuclein SureFire Ultra assay formulations using purified protein standards.
2) Characterization of new total and pS129 α-synuclein SureFire Ultra assay formulations using purified protein standards.
3) Characterization of the most sensitive new total and pS129 α-synuclein SureFire Ultra assay formulations using purified protein standards.
4) Characterization of new total and pS129 α-synuclein SureFire Ultra assays using extracts from mouse brain tissues and HEK293 cells.
5) Evaluation of matrix effects in mouse brain tissue extracts and HEK293 cell lysates using a parallelism experiment.
6) Evaluation of mouse brain tissue matrix effects in total α-synuclein assay using a spike-recovery experiment.
7) Evaluation of mouse brain tissue matrix effects in pS129 α-synuclein assay using a spike-recovery experiment.
8) Sham and PFF mouse brain tissue extract dilutions prior to SureFire Ultra assay measurement.
9) Percentage of total α-synuclein recovered in PBS, Triton and SDS brain tissue fractions from PFF and sham mice.
10) Percentage of pS129 α-synuclein recovered in PBS, Triton and SDS brain tissue fractions from PFF and sham mice.

11) Amount of α-synuclein aggregation quantified in PBS and Triton brain tissue fractions from PFF and sham mice using the LEGEND MAX α-Synuclein Aggregate ELISA.

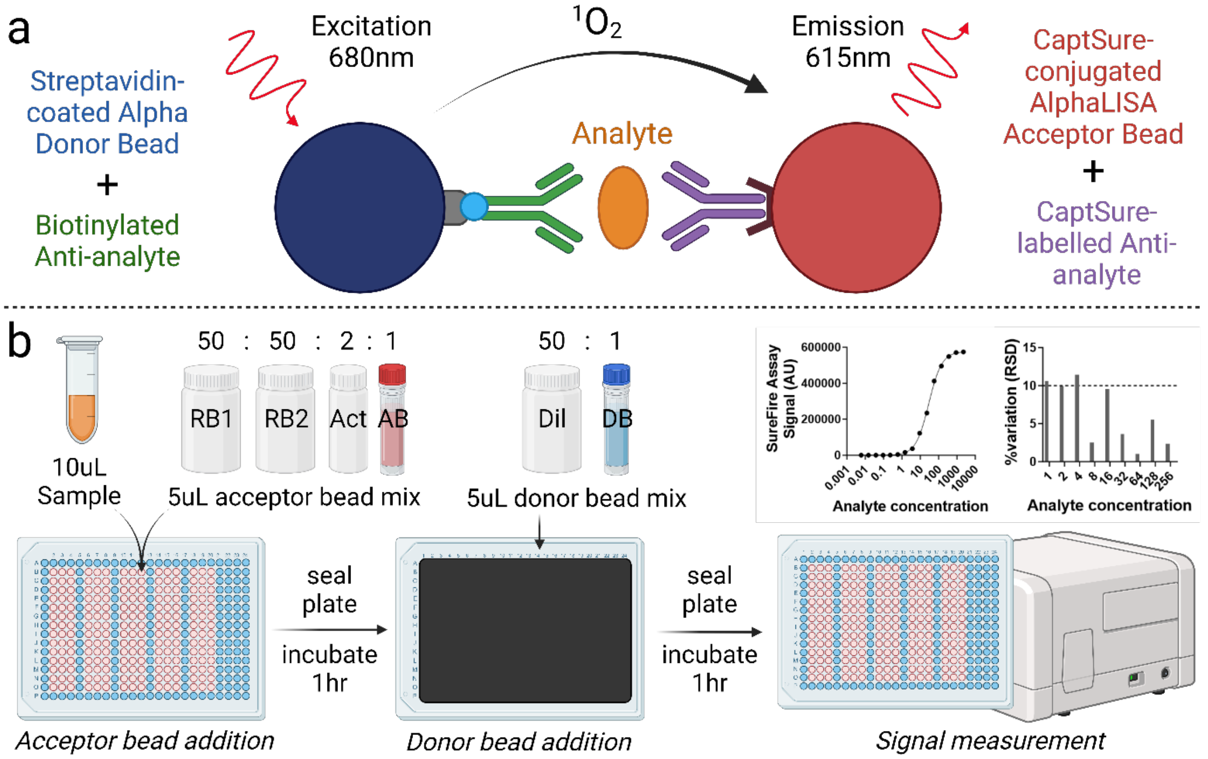


**Supplementary Figure 1. AlphaLISA SureFire Ultra assay platform and workflow schematics.** AlphaLISA SureFire Ultra assays are designed using two antibodies against a target protein, which are differentially tagged to ensure their selective conjugation to one of two types of Alpha beads; donor beads or acceptor beads. The binding of both antibodies to an analyte brings donor and acceptor beads into very close proximity, which enables donor beads to activate acceptor beads following sample photoexcitation to produce the assay signal (**a**). Our assay workflow followed the manufacturer's instructions (**b**). Briefly, reaction buffers 1 (RB1) and 2 (RB2) contain the assay antibodies and are combined with solutions of activation buffer (Act) and acceptor beads (AB) in a 50:50:2:1 ratio, respectively – i.e. 50µL RB1 + 50µL RB2 + 2µL Act + 1µL AB. Five microliters of this solution is added to 10µL of sample in a 384-well microtiter plate (AlphaPlate-384, Perkin Elmer, MA, USA; #6005350), which is then sealed with a light-impermeable microplate seal (TopSeal-A 384, Perkin Elmer; #6050173) and incubated for 1hr at room temperature with orbital rotation (250rpm). Next, donor beads are added to dilution buffer in a 1:50 ratio, before 5µL is added to wells and plates are again sealed and incubated for 1hr at room temperature with orbital rotation. All steps involving donors beads should be performed under subdued laboratory lighting (<100 lux), with green light filters (e.g. LEE090) recommended. Samples were then quantified using an EnVision multimodal plate reader according to manufacturer's AlphaScreen factory default protocol.

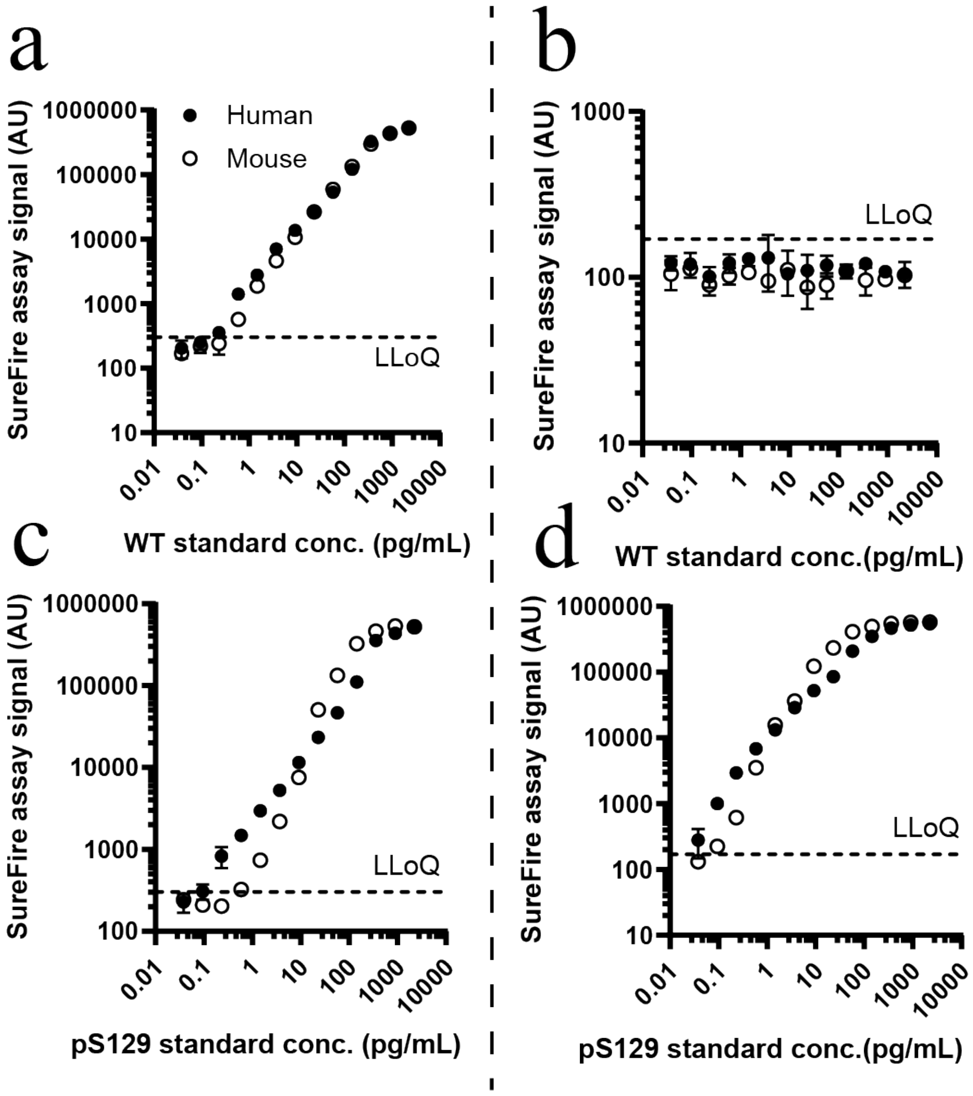


**Supplementary Figure 2. Characterization of the linear dynamic range of new SureFire Ultra assays using purified protein standards.** The linear dynamic range of the most sensitive new formulations of the new total (left; #ALSU-TASYN-B, Revvity) and pS129 (right; #ALSU-PASYN-B, Revvity) α-synuclein assay were characterized using purified human (Hu) and mouse (Ms) wild-type (WT; a-b) and pS129 (c-d) α-synuclein standards. All associated data are contained within Supplementary Tables 2-3. The lower limit of quantification (LLoQ) was defined as 6 standard deviations above the mean of the blank. Data represent mean ± standard deviation. AU, arbitrary units.

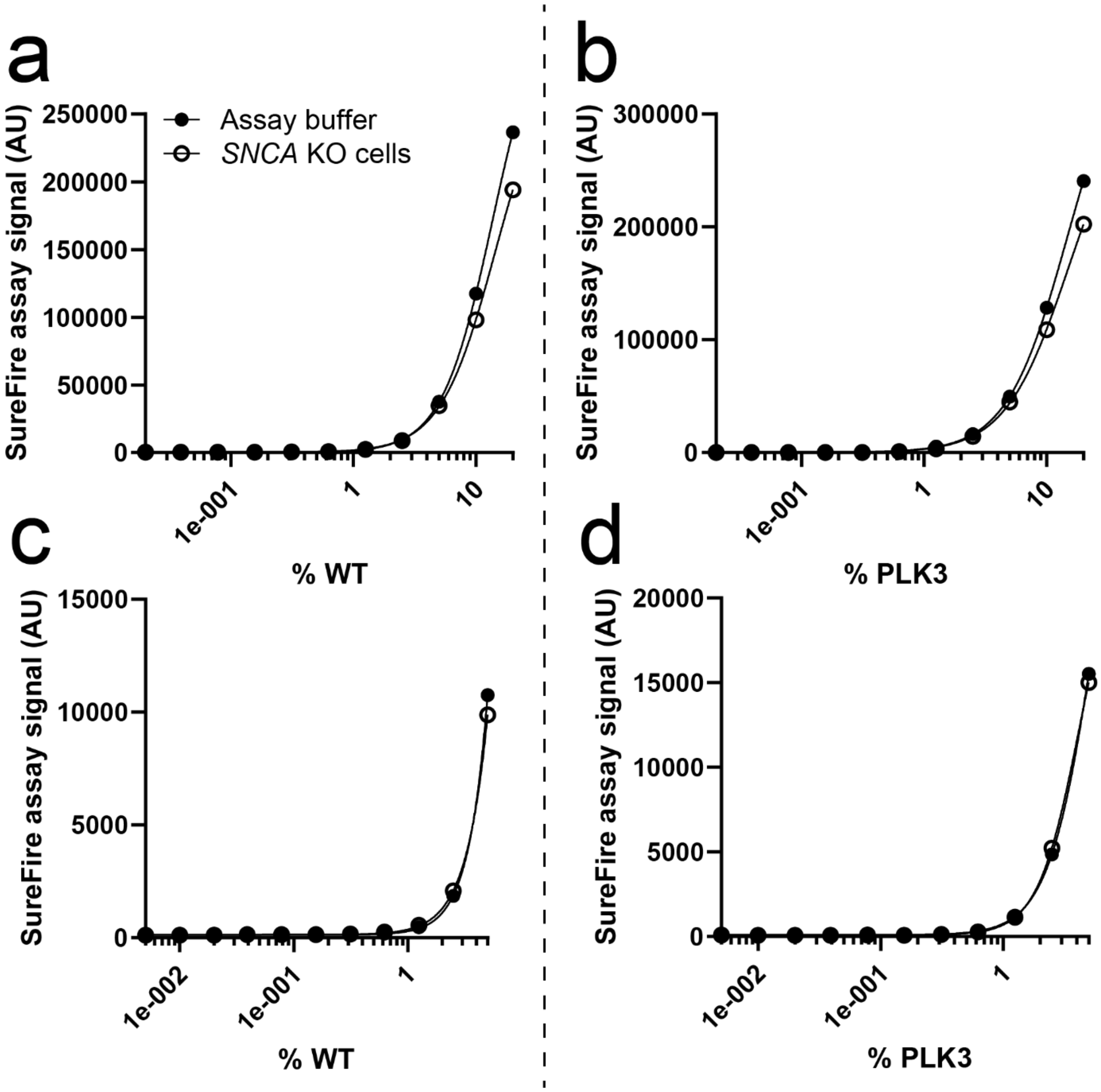


**Supplementary Figure 3. Evaluation of matrix effects in HEK293 cell lysate using a parallelism experiment.** Matrix effects were measured for the most sensitive new total (left; #ALSU-TASYN-B, Revvity) and pS129 (right; #ALSU-PASYN-B, Revvity) α-synuclein assay formulation using 5-fold (**a, b**) and 20-fold (**c, d**) diluted wild-type (WT) and polo-like kinase 3 (PLK3)-transfected HEK293 cell lysates, respectively, diluted in either *SNCA* knock-out (KO) HEK293 cell lysate of an equivalent dilution (5- or 20-fold) or assay buffer. AU, arbitrary units.

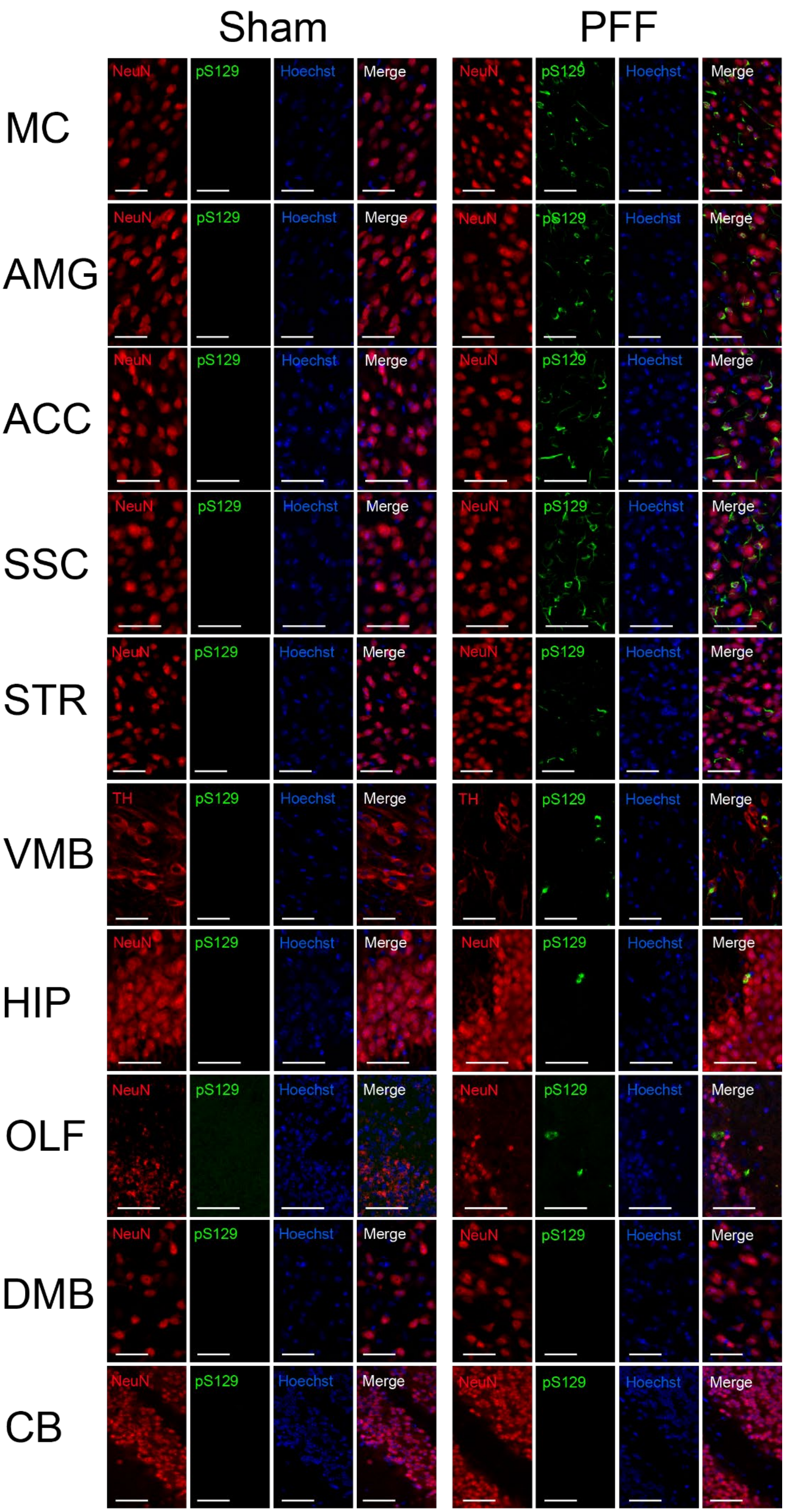

Sham
PFF
MC
AMG
ACC
SSC
STR
VMB
HIP
OLF
DMB
CB
NeuN
pS129
Hoechst
Merge
TH

**Supplementary Figure 4. Immunohistochemical profiles of pS129 α-synuclein across brain regions of interest in sham and PFF mice.** Immunofluorescent labelling of pS129 α-synuclein (EP1536Y, Abcam) was performed in 30µm free-floating, fixed coronal brain tissue sections from 3-month-old PFF and sham mice. Neurons were labelled with neuronal nuclear protein (NeuN; ab279297, Abcam) or tyrosine hydroxylase (TH; ab76442, Abcam) and Hoechst used to counterstain all cell nuclei. Scale bars represent 50µm. ACC, anterior cingulate cortex; AMG, amygdala; CB, cerebellum; DMB, dorso-medial midbrain; HIP, hippocampus; MC, motor cortex; OLF, olfactory bulb; SSC, somatosensory cortex; STR, striatum; VMB, ventral midbrain.

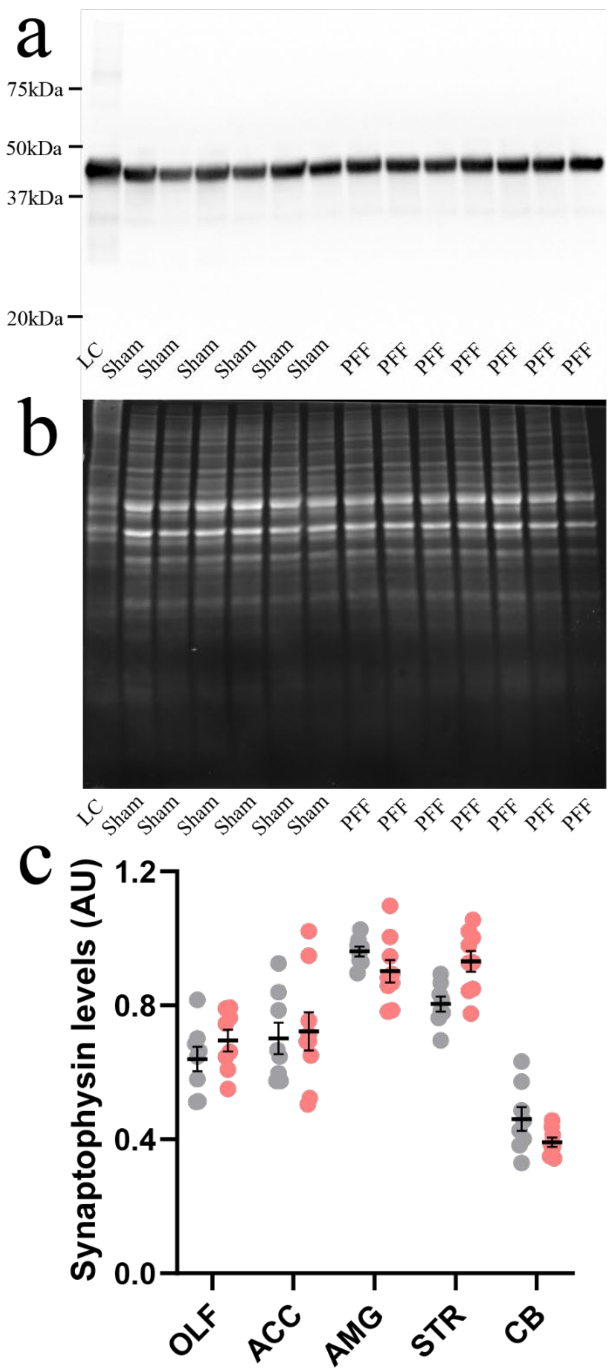


**Supplementary Figure 5. Immunoblot quantification of synaptophysin in sham and PFF mouse brain regions.** Synaptophysin immunoblotting (YE269, Abcam) identified a single synaptophysin isoform (**a**), which was quantified using densitometry and normalized to Sypro Ruby total protein levels to correct against any differences in protein loading between samples (**b**). Representative full blot images above display data from the striatum. Levels of synaptophysin protein were quantified in immunoblots of whole tissue extracts from the olfactory bulb (OLF), anterior cingulate cortex (ACC), amygdala (AMG), striatum (STR) and cerebellum (CB) of sham and PFF mice (**c**). Data represent mean ± standard deviation. AU, arbitrary units. No significant differences were identified between corresponding regions of the sham and PFF mouse brain using a Two-way ANOVA paired with Sidak's multiple comparisons post-hoc tests.

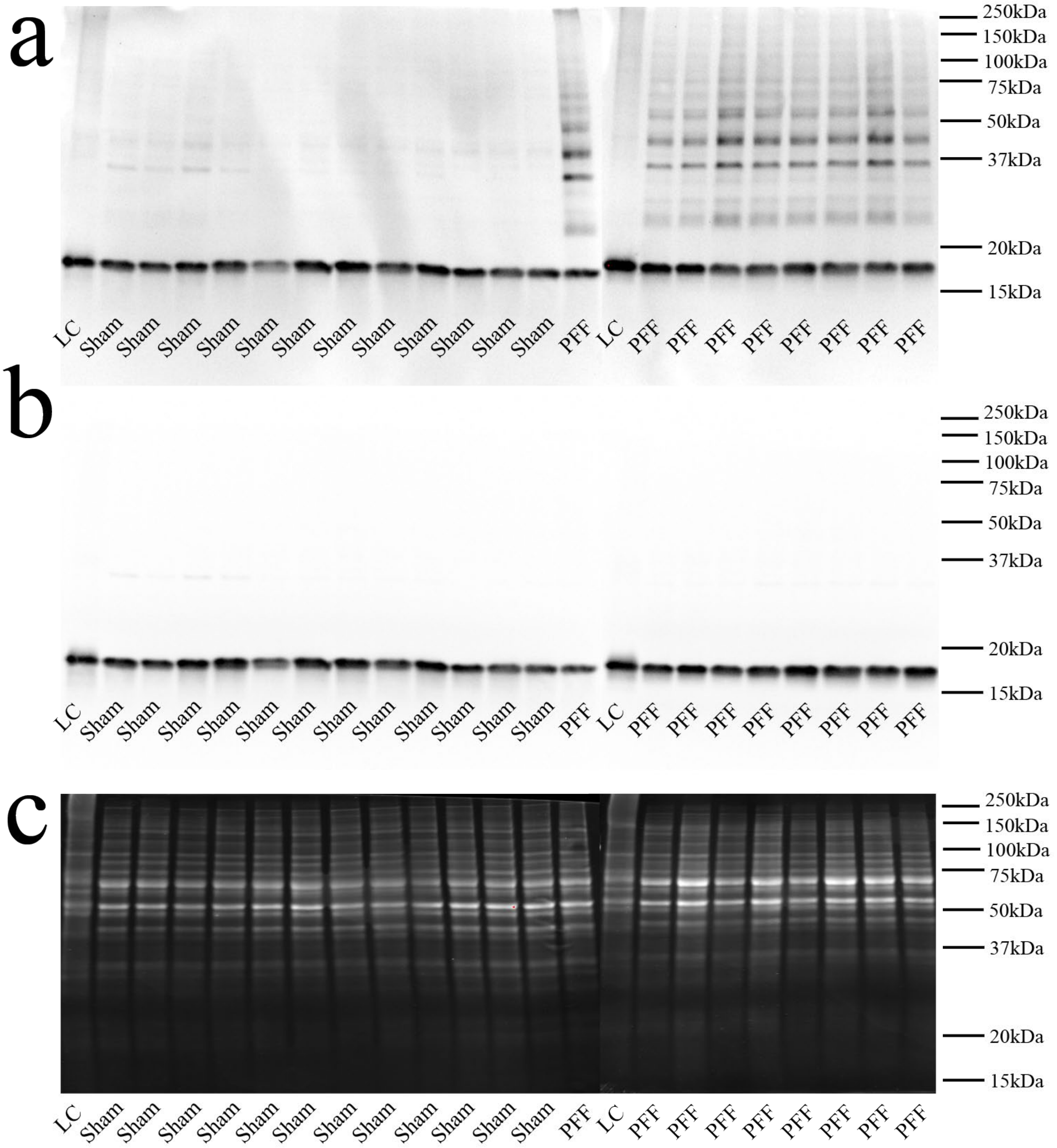


**Supplementary Figure 6. Full representative immunoblots characterising α-synuclein phosphorylation in brain tissue extracts from sham and PFF mice.** α-Synuclein S129 phosphorylation in the anterior cingulate cortex (ACC), striatum and cerebellum of sham and PFF mice was examined using immunoblotting (**a**; D1R1R antibody, Cell Signalling). Immunoblots above are from the ACC only. Membranes were stripped and re-probed for wild-type α-synuclein (**b**; Syn1, BD Biosciences). Wild-type and pS129 α-synuclein band intensities were normalized to Sypro Ruby total protein staining to correct against any differences in protein loading between samples (**c**). A loading control (LC) consisting of pooled sham mouse brain tissue extracts was included on each gel to correct against any differences between immunoblot batches.

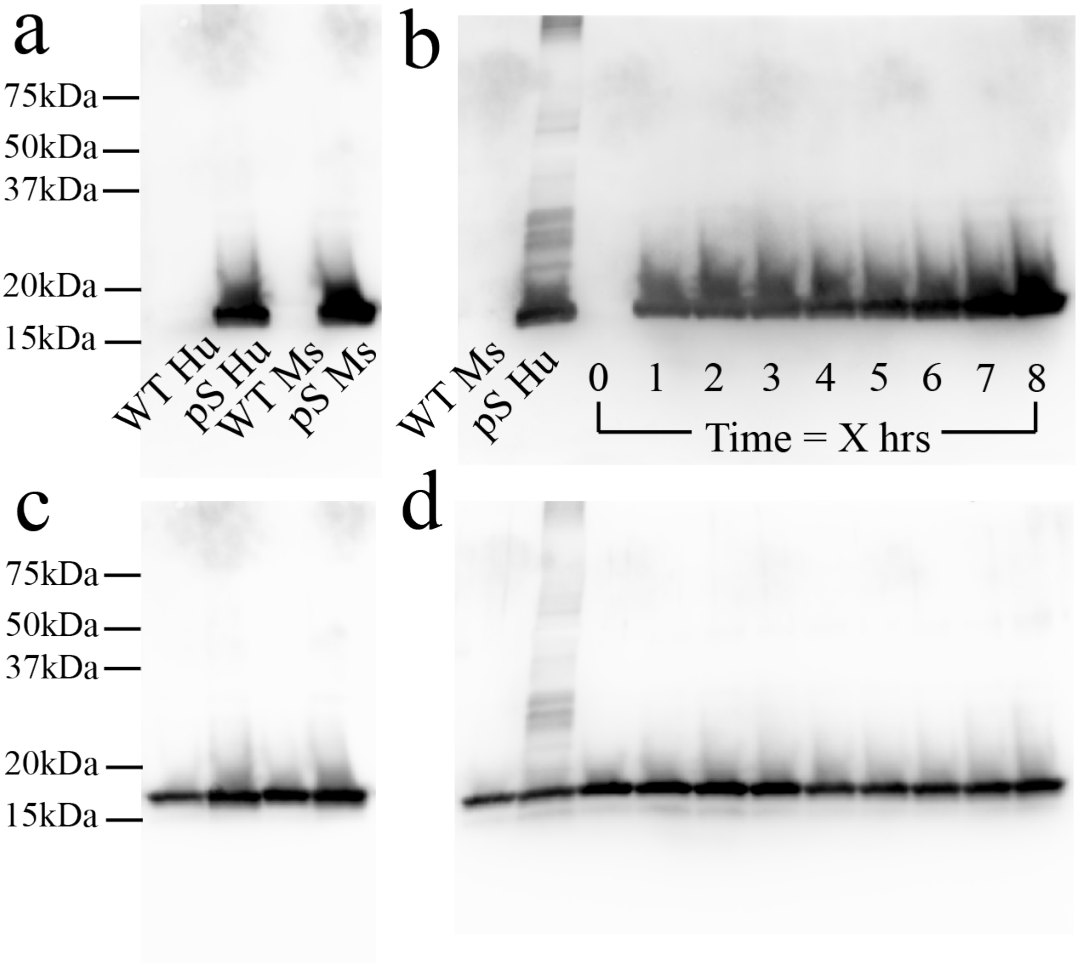


**Supplementary Figure 7. Full representative immunoblots characterising S129 phosphorylation in purified protein standards.** The S129 phosphorylation status of these samples was determined using immunoblotting against human (Hu) and mouse (Ms) pS129 (pS) α-synuclein (D1R1R, Cell Signalling) (**a**). This approach was also used to establish successful phosphorylation of mouse α-synuclein S129 over an 8-hour incubation with human polo-like kinase 3 (PLK3) in kinase reaction buffer (details outlined in methods) (**b**). In both experiments, membranes were stripped and re-probed for human and mouse wild-type (WT) α-synuclein (Syn1, BD Biosciences) to confirm the presence of non-phosphorylated α-synuclein in control lanes (**c, d**), and to demonstrate equal gel loading of α-synuclein over the PLK3 time course experiment (**d**).

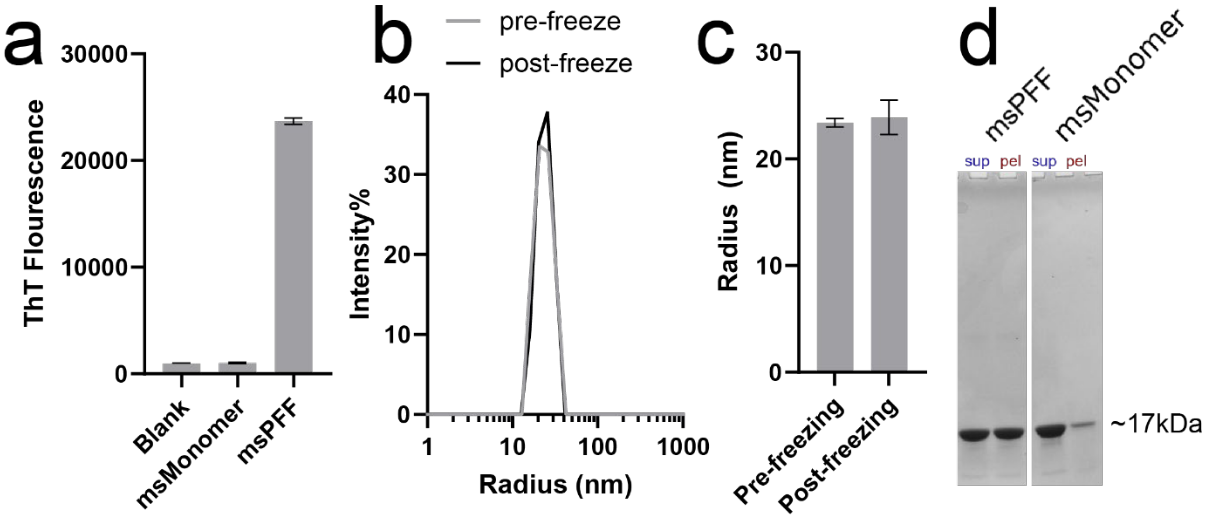


**Supplementary Figure 8. Characterisation of mouse pre-formed fibril (PFF) structure and solubility.** The amyloid structure of mouse α-synuclein PFFs (msPFF) compared with native monomeric mouse α-synuclein (msMonomer) was confirmed using a Thioflavin-T binding assay (**a**). Their fragment size did not change upon freeze-thaw using dynamic light scattering measurements (DynaPro NanoStar, Wyatt Technology) (**b, c**). Sedimentation analysis revealed that approximately 50% of the PFFs became soluble after sonication and freeze/thaw (**d**).

**Supplementary Table 1. Characterization of previous total and pS129 α-synuclein SureFire Ultra assay formulations using purified protein standards.**

| Sample | Protein conc. (pg/mL) | ALSU-TASYN-A total α-synuclein assay (LOD: 156.7, LOQ: 174.4) | | | ALSU-PASYN-A pS129 α-synuclein assay (LOD: 145.7, LOQ: 173.9) | | |
|---|---|---|---|---|---|---|---|
| | | Mean | SD | Inter-assay RSD | Mean | SD | Inter-assay RSD |
| Blank | 0 | 139.0 | 5.9 | 4.2 | 117.5 | 9.4 | 8.0 |
| Mouse pS129 α-synuclein | 0.094 | - | - | - | 135.0 | 9.5 | 7.0 |
| | 0.23 | 135.0 | 10.1 | 7.5 | 123.0 | 7.2 | 5.9 |
| | 0.59 | 145.0 | 12.2 | 8.4 | 144.0 | 4.5 | 3.1 |
| | 1.47 | 139.0 | 21.5 | 15.5 | 158.0 | 10.5 | 6.6 |
| | 3.67 | 135.0 | 9.8 | 7.3 | 359.0 | 33.1 | 9.2 |
| | 9.18 | 137.0 | 20.0 | 14.6 | 594.0 | 40.5 | 6.8 |
| | 22.94 | 148.0 | 23.7 | 16.0 | 1862.0 | 23.0 | 1.2 |
| | 57.34 | 149.0 | 11.1 | 7.4 | 7441.0 | 124.8 | 1.7 |
| | 143.36 | 136.0 | 35.2 | 25.9 | 30567.0 | 1255.2 | 4.1 |
| | 358.4 | 139.0 | 16.3 | 11.7 | 75618.0 | 998.4 | 1.3 |
| | 896 | 142.0 | 10.6 | 7.5 | 175823.0 | 11153.1 | 6.3 |
| | 2240 | 147.0 | 10.6 | 7.2 | 310631.0 | 5396.7 | 1.7 |
| | 5600 | 156.0 | 5.2 | 3.3 | 356138.0 | 16296.8 | 4.6 |
| | 14000 | 153.0 | 4.6 | 3.0 | - | - | - |
| | 28000 | 167.0 | 19.0 | 11.4 | - | - | - |
| | 56000 | 175.0 | 3.8 | 2.2 | - | - | - |
| Mouse WT α-synuclein | 0.094 | - | - | - | 119 | 7.5 | 6.3 |
| | 0.23 | 156 | 4.4 | 2.8 | 135 | 8.3 | 6.1 |
| | 0.59 | 167 | 2.3 | 1.4 | 137 | 12.8 | 9.3 |
| | 1.47 | 172 | 6.7 | 3.9 | 148 | 26.1 | 17.6 |
| | 3.67 | 179 | 6.1 | 3.4 | 129 | 11.2 | 8.7 |
| | 9.18 | 172 | 2.9 | 1.7 | 136 | 4.5 | 3.3 |
| | 22.94 | 172 | 6.7 | 3.9 | 139 | 4.6 | 3.3 |
| | 57.34 | 175 | 8.9 | 5.1 | 122 | 11 | 9.0 |
| | 143.36 | 161 | 4 | 2.5 | 147 | 9.8 | 6.7 |
| | 358.4 | 176 | 6.7 | 3.8 | 116 | 13.2 | 11.4 |
| | 896 | 210 | 26.7 | 12.7 | 133 | 41.2 | 31.0 |
| | 2240 | 280 | 4 | 1.4 | 137.0 | 25.6 | 18.7 |
| | 5600 | 293.0 | 2.1 | 0.7 | 175.0 | 43.7 | 25.0 |
| | 14000 | 647.0 | 67.0 | 10.4 | - | - | - |
| | 28000 | 1414.0 | 98.4 | 7.0 | - | - | - |
| | 56000 | 3263.0 | 91.7 | 2.8 | - | - | - |
| Human pS129 α-synuclein | 0.094 | - | - | - | 149 | 9.9 | 6.6 |
| | 0.23 | 119.0 | 4.0 | 3.4 | 110 | 12.1 | 11.0 |
| | 0.59 | 132.0 | 5.2 | 3.9 | 123 | 10.5 | 8.5 |
| | 1.47 | 187.0 | 6.9 | 3.7 | 125 | 8.5 | 6.8 |
| | 3.67 | 242.0 | 8.9 | 3.7 | 115 | 12.5 | 10.9 |
| | 9.18 | 734.0 | 16.0 | 2.2 | 105 | 11.2 | 10.7 |
| | 22.94 | 2209.0 | 61.7 | 2.8 | 129 | 13.2 | 10.2 |
| | 57.34 | 3866.0 | 150.7 | 3.9 | 123 | 11.2 | 9.1 |
| | 143.36 | 9648.0 | 820.8 | 8.5 | 126 | 0.6 | 0.5 |
| | 358.4 | 21824.0 | 1569.6 | 7.2 | 99 | 8.7 | 8.8 |
| | 896 | 85624.0 | 5052.7 | 5.9 | 110 | 4.6 | 4.2 |
| | 2240 | 187110.0 | 10976.3 | 5.9 | 102.0 | 10.5 | 10.3 |
| | 5600 | 318475.0 | 25955.1 | 8.1 | 511.0 | 36.2 | 7.1 |
| | 14000 | 360226.0 | 29371.4 | 8.2 | - | - | - |
| | 28000 | 368963.0 | 15569.5 | 4.2 | - | - | - |

| | | | | | | | |
|---|---|---|---|---|---|---|---|
| | 56000 | 370536.0 | 17555.3 | 4.7 | - | - | - |
| Human WT α-synuclein | 0.094 | - | - | - | 112 | 3.6 | 3.2 |
| | 0.23 | 124.0 | 3.0 | 2.4 | 121 | 2.3 | 1.9 |
| | 0.59 | 134.0 | 4.5 | 3.4 | 147 | 5.2 | 3.5 |
| | 1.47 | 144.0 | 7.1 | 4.9 | 172 | 11.1 | 6.5 |
| | 3.67 | 173.0 | 14.0 | 8.1 | 113 | 20.9 | 18.5 |
| | 9.18 | 224.0 | 10.0 | 4.5 | 116 | 12.1 | 10.4 |
| | 22.94 | 321.0 | 33.3 | 10.4 | 120 | 19.3 | 16.1 |
| | 57.34 | 640.0 | 52.8 | 8.3 | 145 | 10.6 | 7.3 |
| | 143.36 | 1278.0 | 312.6 | 24.5 | 120 | 10.6 | 8.8 |
| | 358.4 | 2308.0 | 420.8 | 18.2 | 123 | 17.2 | 14.0 |
| | 896 | 5836.0 | 1110.5 | 19.0 | 123 | 8.9 | 7.2 |
| | 2240 | 22559.0 | 2050.0 | 9.1 | 120.0 | 8.9 | 7.4 |
| | 5600 | 85978.0 | 8426.3 | 9.8 | 145.0 | 9.8 | 6.8 |
| | 14000 | 234311.0 | 15926.1 | 6.8 | - | - | - |
| | 28000 | 336622.0 | 26371.3 | 7.8 | - | - | - |
| | 56000 | 359868.0 | 19563.1 | 5.4 | - | - | - |

Existing formulations of the total (#ALSU-TASYN-A, Revvity) and pS129 (#ALSU-PASYN-A, Revvity) α-synuclein assay were characterized using purified human and mouse wild-type (WT) and pS129 α-synuclein standards. Inter-assay variation (RSD) was defined as the standard deviation expressed as a percentage of the mean. The limit of detection (LoD) and lower limit of quantification (LLoQ) were defined as 3 and 6 standard deviations above the mean of the blank, respectively. Data represent raw assay signal measurements in arbitrary units.

**Supplementary Table 2. Characterization of new total and pS129 α-synuclein SureFire Ultra assay formulations using purified protein standards.**

| α-syn std | Standard conc. (ng/mL) | New total α-synuclein assay formulations | | | | | New pS129 α-synuclein assay formulations | | | | |
|---|---|---|---|---|---|---|---|---|---|---|---|
| | | 1 | 2 | 3 | 4 | 5 | 1 | 2 | 3 | 4 | 5 |
| Mouse p-S129 | 20 | 840792 | 1029656 | 742576 | 341 | 1593 | 998587 | 756485 | 1067388 | 938 | 486616 |
| | | 910771 | 954597 | 693577 | 290 | 1391 | 939358 | 734843 | 964689 | 975 | 479148 |
| | 2 | 277935 | 194720 | 129172 | 197 | 743 | 327756 | 193746 | 481042 | 415 | 75927 |
| | | 285257 | 194973 | 129277 | 231 | 559 | 326086 | 194712 | 483272 | 378 | 73662 |
| | 0.2 | 6079 | 1891 | 4855 | 205 | 244 | 22037 | 17900 | 29594 | 222 | 4674 |
| | | 7006 | 1326 | 4759 | 172 | 243 | 20228 | 16816 | 28332 | 214 | 4401 |
| | 0 | 303 | 273 | 306 | 210 | 180 | 281 | 433 | 324 | 210 | 243 |
| | | 323 | 256 | 276 | 193 | 235 | 176 | 240 | 219 | 193 | 184 |
| Mouse WT | 20 | 657404 | 742985 | 480785 | 201 | 205 | 218 | 290 | 252 | 151 | 201 |
| | | 644834 | 739840 | 482842 | 256 | 168 | 226 | 260 | 210 | 201 | 252 |
| | 2 | 94995 | 25682 | 42098 | 240 | 159 | 235 | 218 | 189 | 197 | 201 |
| | | 95535 | 21375 | 38472 | 264 | 189 | 189 | 243 | 189 | 168 | 180 |
| | 0.2 | 2303 | 595 | 2286 | 197 | 168 | 213 | 264 | 193 | 180 | 243 |
| | | 1770 | 518 | 2343 | 227 | 202 | 201 | 239 | 227 | 184 | 168 |
| | 0 | 214 | 227 | 277 | 210 | 176 | 210 | 268 | 210 | 168 | 184 |
| | | 218 | 243 | 239 | 277 | 197 | 193 | 222 | 210 | 205 | 243 |
| Human p-S129 | 20 | 1340 | 811240 | 665 | 175812 | 849430 | 1165 | 576515 | 881121 | 692920 | 465942 |
| | | 1242 | 824176 | 590 | 190495 | 872589 | 1158 | 595343 | 871329 | 675019 | 462341 |
| | 2 | 317 | 67561 | 271 | 2762 | 134524 | 405 | 112253 | 347305 | 175893 | 72925 |
| | | 251 | 62621 | 267 | 2184 | 136104 | 326 | 103457 | 301306 | 164545 | 67799 |
| | 0.2 | 196 | 1168 | 268 | 254 | 3117 | 246 | 12435 | 27363 | 19077 | 8347 |
| | | 188 | 1093 | 225 | 288 | 2676 | 225 | 13363 | 30153 | 22601 | 9179 |
| | 0 | 213 | 225 | 230 | 213 | 196 | 196 | 263 | 267 | 259 | 221 |
| | | 376 | 228 | 284 | 841 | 200 | 221 | 238 | 180 | 188 | 205 |
| Human WT | 20 | 1474 | 663827 | 523 | 134131 | 771906 | 301 | 247 | 230 | 230 | 221 |
| | | 1485 | 670452 | 552 | 132822 | 722961 | 334 | 263 | 188 | 225 | 196 |
| | 2 | 306 | 22796 | 280 | 1676 | 57367 | 234 | 221 | 213 | 184 | 217 |
| | | 268 | 20476 | 259 | 1337 | 52731 | 209 | 213 | 209 | 200 | 234 |
| | 0.2 | 222 | 427 | 242 | 255 | 788 | 192 | 234 | 188 | 188 | 175 |
| | | 188 | 373 | 272 | 188 | 683 | 209 | 200 | 201 | 171 | 180 |
| | 0 | 217 | 276 | 268 | 205 | 213 | 196 | 406 | 205 | 184 | 225 |

| | | | | | | | | | | | |
|---|---|---|---|---|---|---|---|---|---|---|---|
| | | 213 | 264 | 242 | 238 | 201 | 222 | 234 | 205 | 201 | 196 |

Five new formulations of the total and pS129 α-synuclein assay were characterized using purified human and mouse wild-type (WT) and pS129 α-synuclein standards. Data represent mean raw assay signal measurements in arbitrary units.

**Supplementary Table 3. Characterization of the most sensitive new total and pS129 α-synuclein SureFire Ultra assay formulations using purified protein standards.**

| Sample | Protein conc. (pg/mL) | New Total α-synuclein pairing 2 (LOD: 233.1, LOQ: 304.2) | | | New pS129 α-synuclein pairing 3 (LOD: 150.7, LOQ: 192.1) | | |
|---|---|---|---|---|---|---|---|
| | | Mean | SD | Inter-assay RSD | Mean | SD | Inter-assay RSD |
| Blank | 0 | 162.0 | 23.7 | 14.6 | 109.3 | 13.8 | 12.6 |
| Mouse pS129 α-synuclein | 0.038 | 200.0 | 13.3 | 6.7 | 132.0 | 2.0 | 1.5 |
| | 0.094 | 203.0 | 25.1 | 12.4 | 224.0 | 7.8 | 3.5 |
| | 0.23 | 325.0 | 24.7 | 7.6 | 617.0 | 58.7 | 9.5 |
| | 0.59 | 640.0 | 30.4 | 4.8 | 3523.0 | 97.5 | 2.8 |
| | 1.47 | 2190.0 | 57.9 | 2.6 | 15815.0 | 176.1 | 1.1 |
| | 3.67 | 7583.0 | 363.9 | 4.8 | 36411.0 | 919.1 | 2.5 |
| | 9.18 | 50570.0 | 3329.9 | 6.6 | 122497.0 | 1745.1 | 1.4 |
| | 22.94 | 133627.0 | 12612.2 | 9.4 | 234623.0 | 3148.9 | 1.3 |
| | 57.34 | 323984.0 | 10499.0 | 3.2 | 412329.0 | 13342.7 | 3.2 |
| | 143.36 | 462248.0 | 7490.4 | 1.6 | 495793.0 | 3648.9 | 0.7 |
| | 358.4 | 533905.0 | 16465.8 | 3.1 | 548751.0 | 38523.1 | 7.0 |
| | 896 | 524503.0 | 37267.9 | 7.1 | 570531.0 | 22952.3 | 4.0 |
| | 2240 | 543322.0 | 47032.3 | 8.7 | 574021.0 | 26438.8 | 4.6 |
| Mouse WT α-synuclein | 0.038 | 169 | 13.7 | 8.1 | 105 | 21.5 | 20.5 |
| | 0.094 | 219 | 46.4 | 21.2 | 113 | 5 | 4.4 |
| | 0.23 | 241 | 79 | 32.8 | 90 | 12.1 | 13.4 |
| | 0.59 | 568 | 9.3 | 1.6 | 102 | 11.6 | 11.4 |
| | 1.47 | 1869 | 123.5 | 6.6 | 107 | 9.3 | 8.7 |
| | 3.67 | 4619 | 405.5 | 8.8 | 95 | 8.1 | 8.5 |
| | 9.18 | 10629 | 926.1 | 8.7 | 111 | 33.6 | 30.3 |
| | 22.94 | 26534 | 1910.8 | 7.2 | 87 | 22.6 | 26.0 |
| | 57.34 | 59111 | 4623.2 | 7.8 | 90 | 15.6 | 17.3 |
| | 143.36 | 134113 | 11574.5 | 8.6 | 109 | 9.3 | 8.5 |
| | 358.4 | 298003 | 17099.7 | 5.7 | 96 | 18.2 | 19.0 |
| | 896 | 435612 | 23564 | 5.4 | 97 | 5.2 | 5.4 |
| | 2240 | 526356 | 35263 | 6.7 | 103 | 4.5 | 4.4 |
| Human pS129 α-synuclein | 0.038 | 228 | 59.4 | 26.1 | 280 | 132.5 | 47.3 |
| | 0.094 | 308 | 65.6 | 21.3 | 1004 | 28.3 | 2.8 |
| | 0.23 | 831 | 239.4 | 28.8 | 2935 | 108.5 | 3.7 |
| | 0.59 | 1489 | 120.5 | 8.1 | 6834 | 367.8 | 5.4 |
| | 1.47 | 2960 | 486.1 | 16.4 | 13260 | 1939.7 | 14.6 |
| | 3.67 | 5270 | 518.5 | 9.8 | 28740 | 2849.7 | 9.9 |
| | 9.18 | 11565 | 461 | 4.0 | 52311 | 7528.7 | 14.4 |
| | 22.94 | 23302 | 2141 | 9.2 | 85369 | 2123.3 | 2.5 |
| | 57.34 | 46691 | 1640.3 | 3.5 | 208784 | 30487 | 14.6 |
| | 143.36 | 111313 | 11016 | 9.9 | 350602 | 12652.2 | 3.6 |
| | 358.4 | 356486 | 19709.8 | 5.5 | 468104 | 4513 | 1.0 |
| | 896 | 434334 | 56644 | 13.0 | 516624 | 28545.8 | 5.5 |
| | 2240 | 518737 | 32622.4 | 6.3 | 545949 | 13291.6 | 2.4 |
| Human WT α-synuclein | 0.038 | 205 | 63.3 | 30.9 | 122 | 12.1 | 9.9 |
| | 0.094 | 253 | 59 | 23.3 | 120 | 20.6 | 17.2 |
| | 0.23 | 357 | 12.1 | 3.4 | 101 | 14.4 | 14.3 |
| | 0.59 | 1403 | 212.3 | 15.1 | 122 | 15.6 | 12.8 |
| | 1.47 | 2782 | 235.7 | 8.5 | 129 | 12.1 | 9.4 |
| | 3.67 | 7048 | 684.5 | 9.7 | 131 | 49 | 37.4 |
| | 9.18 | 13682 | 1689.8 | 12.4 | 105 | 7.1 | 6.8 |
| | 22.94 | 25904 | 2414.4 | 9.3 | 110 | 26.7 | 24.3 |

| | | | | | | | |
|---|---|---|---|---|---|---|---|
| | 57.34 | 53230 | 5042.2 | 9.5 | 118 | 16.9 | 14.3 |
| | 143.36 | 120243 | 15425.4 | 12.8 | 109 | 10.4 | 9.5 |
| | 358.4 | 326893 | 15281.9 | 4.7 | 121 | 4.5 | 3.7 |
| | 896 | 449586 | 23526 | 5.2 | 108 | 4.5 | 4.2 |
| | 2240 | 522986 | 29653.2 | 5.7 | 105 | 18.7 | 17.8 |

The most sensitive new total (#ALSU-TASYN-B, Revvity) and pS129 (#ALSU-PASYN-B, Revvity) α-synuclein assay formulation were characterized using purified human and mouse wild-type (WT) and pS129 α-synuclein standards. Inter-assay variation (RSD) was defined as the standard deviation expressed as a percentage of the mean. The limit of detection (LoD) and lower limit of quantification (LLoQ) were defined as 3 and 6 standard deviations above the mean of the blank, respectively. Data represent raw assay signal measurements in arbitrary units.

**Supplementary Table 4. Characterization of new total and pS129 α-synuclein SureFire Ultra assays using extracts from mouse brain tissues and HEK293 cells.**

| Sample | Dilution factor | New Total α-synuclein pairing 2 (LOD: 233.1, LOQ: 304.2) | | | New total α-synuclein pairing 3 (LOD: 259.4, LOQ: 391.1) | | | New pS129 α-synuclein pairing 3 (LOD: 150.7, LOQ: 192.1) | | | New pS129 α-synuclein pairing 2 (LOD: 141.2, LOQ: 179.5) | | |
|---|---|---|---|---|---|---|---|---|---|---|---|---|---|
| | | Mean | SD | Inter-assay RSD | Mean | SD | Inter-assay RSD | Mean | SD | Inter-assay RSD | Mean | SD | Inter-assay RSD |
| WT mouse brain tissue | Neat | 519383.0 | 20653.2 | 4.0 | 477170.0 | 18897.9 | 4.0 | 128664.0 | 1074.7 | 0.8 | 31413.0 | 2453.8 | 7.8 |
| | 2 | 527502.0 | 6619.0 | 1.3 | 491094.0 | 4284.9 | 0.9 | 126258.0 | 8867.9 | 7.0 | 31082.0 | 5546.3 | 17.8 |
| | 4 | 529393.0 | 29850.5 | 5.6 | 486487.0 | 23452.7 | 4.8 | 124355.0 | 10461.7 | 8.4 | 31520.0 | 5469.4 | 17.4 |
| | 8 | 525027.0 | 49416.3 | 9.4 | 484084.0 | 8826.4 | 1.8 | 119105.0 | 3926.0 | 3.3 | 28206.0 | 2488.5 | 8.8 |
| | 16 | 528122.0 | 13042.0 | 2.5 | 477008.0 | 11361.4 | 2.4 | 103232.0 | 6748.2 | 6.5 | 29853.0 | 2577.5 | 8.6 |
| | 32 | 515598.0 | 7265.3 | 1.4 | 486933.0 | 35996.6 | 7.4 | 72292.0 | 1043.0 | 1.4 | 30285.0 | 1248.8 | 4.1 |
| | 64 | 518405.0 | 12465.6 | 2.4 | 453018.0 | 21384.7 | 4.7 | 37331.0 | 878.8 | 2.4 | 19466.0 | 1837.6 | 9.4 |
| | 128 | 504747.0 | 14532.4 | 2.9 | 409767.0 | 7796.9 | 1.9 | 15619.0 | 616.8 | 3.9 | 11628.0 | 569.8 | 4.9 |
| | 256 | 485607.0 | 15069.2 | 3.1 | 338408.0 | 16359.1 | 4.8 | 5091.0 | 88.6 | 1.7 | 6000.0 | 459.1 | 7.7 |
| | 512 | 422613.0 | 6398.5 | 1.5 | 241884.0 | 324.8 | 0.1 | 1561.0 | 90.6 | 5.8 | 3119.0 | 158.6 | 5.1 |
| | 1024 | 284070.0 | 17109.5 | 6.0 | 146147.0 | 6118.5 | 4.2 | 538.0 | 42.4 | 7.9 | 1552.0 | 65.5 | 4.2 |
| | 2048 | 132522.0 | 6133.0 | 4.6 | 70906.0 | 1376.3 | 1.9 | 237.0 | 14.3 | 6.0 | 806.0 | 66.9 | 8.3 |
| | 4096 | 49527.0 | 1159.6 | 2.3 | 36135.0 | 749.9 | 2.1 | 148.0 | 5.8 | 3.9 | 515.0 | 19.0 | 3.7 |
| | 8192 | 15882.0 | 1754.0 | 11.0 | 15976.0 | 439.9 | 2.8 | 107.0 | 2.9 | 2.7 | 356.0 | 13.7 | 3.8 |
| | 16384 | 3286.0 | 96.4 | 2.9 | 5876.0 | 211.6 | 3.6 | 101.0 | 11.1 | 11.0 | 248.0 | 29.5 | 11.9 |
| | 32768 | 725.0 | 56.1 | 7.7 | 1919.0 | 40.0 | 2.1 | 91.0 | 16.2 | 17.8 | 177.0 | 31.8 | 18.0 |
| | Blank | 126 | 9.0 | 7.1 | 142 | 24.9 | 17.5 | 90 | 12.1 | 13.5 | 105 | 10.4 | 9.9 |
| Snca-/- mouse brain tissue | Neat | 1789 | 147.2 | 8.2 | 1940 | 239.1 | 12.3 | 89 | 4.5 | 5.1 | 97 | 5.2 | 5.4 |
| | 2 | 1332 | 60.4 | 4.5 | 1209 | 107.5 | 8.9 | 100 | 25.9 | 25.9 | 88 | 5.2 | 5.9 |
| | 4 | 755 | 120 | 15.9 | 1143 | 100.6 | 8.8 | 70 | 13.7 | 19.6 | 95 | 10.4 | 10.9 |
| | 8 | 406 | 67 | 16.5 | 920 | 142.5 | 15.5 | 76 | 13.5 | 17.8 | 112 | 15.9 | 14.2 |
| | 16 | 363 | 32.3 | 8.9 | 805 | 160.5 | 19.9 | 103 | 4.5 | 4.4 | 107 | 19.2 | 17.9 |
| | 32 | 238 | 26.1 | 11.0 | 677 | 130.5 | 19.3 | 116 | 47.1 | 40.6 | 119 | 13.7 | 11.5 |
| | 64 | 189 | 22.6 | 12.0 | 554 | 73.6 | 13.3 | 108 | 22.1 | 20.5 | 114 | 25.2 | 22.1 |
| | 128 | 146 | 13.7 | 9.4 | 355 | 67.5 | 19.0 | 110 | 13 | 11.8 | 120 | 15.6 | 13.0 |
| | 256 | 122 | 14.4 | 11.8 | 247 | 31.6 | 12.8 | 113 | 27 | 23.9 | 129 | 18 | 14.0 |
| | 512 | 120 | 22 | 18.3 | 201 | 7.8 | 3.9 | 92 | 5.2 | 5.7 | 99 | 11.1 | 11.2 |

| | | | | | | | | | | | | | |
|---|---|---|---|---|---|---|---|---|---|---|---|---|---|
| | 1024 | 98 | 9 | 9.2 | 153 | 27.5 | 18.0 | 113 | 6.7 | 5.9 | 99 | 13.7 | 13.8 |
| | 2048 | 111 | 18 | 16.2 | 144 | 8.9 | 6.2 | 96 | 18.2 | 19.0 | 122 | 12.7 | 10.4 |
| | 4096 | 126 | 8.9 | 7.1 | 140 | 11.6 | 8.3 | 96 | 20.4 | 21.3 | 113 | 10.7 | 9.5 |
| | Blank | 121 | 10.1 | 8.3 | 126 | 18.7 | 14.9 | 93 | 9.0 | 9.7 | 108 | 6.6 | 6.1 |
| WT HEK293 cells | Neat | 458592 | 16881.9 | 3.7 | 4531 | 488.6 | 10.8 | 10856 | 504.9 | 4.7 | 7946 | 149 | 1.9 |
| | 2 | 467632 | 3642.1 | 0.8 | 2243 | 501 | 22.3 | 6407 | 958.2 | 15.0 | 6821 | 222.7 | 3.3 |
| | 4 | 440610 | 15093.4 | 3.4 | 720 | 62.4 | 8.7 | 2621 | 68.5 | 2.6 | 3863 | 211.2 | 5.5 |
| | 8 | 371134 | 12992.4 | 3.5 | 304 | 23.1 | 7.6 | 865 | 146.3 | 16.9 | 2029 | 76.6 | 3.8 |
| | 16 | 266379 | 10172.9 | 3.8 | 222 | 16.5 | 7.4 | 313 | 18.8 | 6.0 | 1220 | 23.3 | 1.9 |
| | 32 | 130916 | 6764.5 | 5.2 | 155 | 18.2 | 11.7 | 172 | 36.9 | 21.5 | 690 | 17 | 2.5 |
| | 64 | 46238 | 3589.1 | 7.8 | 162 | 4.5 | 2.8 | 128 | 13.3 | 10.4 | 411 | 38.2 | 9.3 |
| | 128 | 9315 | 1828.2 | 19.6 | 170 | 2.6 | 1.5 | 142 | 21 | 14.8 | 281 | 16.5 | 5.9 |
| | 256 | 2184 | 184.1 | 8.4 | 165 | 14.3 | 8.7 | 127 | 21.5 | 16.9 | 236 | 9.5 | 4.0 |
| | 512 | 607 | 41.7 | 6.9 | 155 | 17.2 | 11.1 | 117 | 13.5 | 11.5 | 162 | 7.5 | 4.6 |
| | 1024 | 313 | 73.7 | 23.5 | 174 | 9.3 | 5.3 | 105 | 11.6 | 11.0 | 144 | 20.8 | 14.4 |
| | 2048 | 230 | 31.2 | 13.6 | 189 | 25.3 | 13.4 | 130 | 27.2 | 20.9 | 141 | 6.7 | 4.8 |
| | 4096 | 203 | 11.7 | 5.8 | 180 | 29.7 | 16.5 | 106 | 13.7 | 12.9 | 132 | 14.4 | 10.9 |
| | Blank | 169 | 12.7 | 7.5 | 133 | 9.5 | 7.1 | 91 | 14.6 | 16.0 | 127 | 10.4 | 8.2 |
| PLK3-transfected HEK293 cells | Neat | 342392 | 9226.1 | 2.7 | 2531 | 350 | 13.8 | 150754 | 6274.2 | 4.2 | 120563 | 7985.1 | 6.6 |
| | 2 | 327276 | 2669.2 | 0.8 | 2051 | 600 | 29.3 | 127962 | 2422 | 1.9 | 100548 | 3564.5 | 3.5 |
| | 4 | 276227 | 2739.1 | 1.0 | 532 | 55 | 10.3 | 66400 | 1044.1 | 1.6 | 56213 | 2563.2 | 4.6 |
| | 8 | 225865 | 1241.1 | 0.5 | 295 | 15 | 5.1 | 26010 | 1297.2 | 5.0 | 20356 | 1985.6 | 9.8 |
| | 16 | 148819 | 4469.7 | 3.0 | 275 | 26 | 9.5 | 8887 | 392.6 | 4.4 | 6594 | 562.4 | 8.5 |
| | 32 | 62637 | 6088.7 | 9.7 | 154 | 28 | 18.2 | 2413 | 169.8 | 7.0 | 2256 | 156.9 | 7.0 |
| | 64 | 18720 | 484 | 2.6 | 165 | 5.1 | 3.1 | 756 | 42.2 | 5.6 | 859 | 56.9 | 6.6 |
| | 128 | 4274 | 303.7 | 7.1 | 162 | 3.2 | 2.0 | 261 | 13.3 | 5.1 | 302 | 5.2 | 1.7 |
| | 256 | 991 | 84.6 | 8.5 | 166 | 5.3 | 3.2 | 130 | 10.4 | 8.0 | 152 | 15.2 | 10.0 |
| | 512 | 317 | 11.8 | 3.7 | 156 | 6.2 | 4.0 | 114 | 7.1 | 6.2 | 140 | 11.2 | 8.0 |
| | 1024 | 166 | 14.7 | 8.9 | 153 | 19.2 | 12.5 | 115 | 24.9 | 21.7 | 141 | 25 | 17.7 |
| | 2048 | 127 | 10.8 | 8.5 | 165 | 2.1 | 1.3 | 90 | 11.6 | 12.9 | 135 | 16.2 | 12.0 |
| | 4096 | 123 | 16.3 | 13.3 | 168 | 7.5 | 4.5 | 95 | 9.8 | 10.3 | 140 | 12.3 | 8.8 |
| | Blank | 122 | 19.1 | 15.6 | 125 | 21.0 | 16.8 | 73 | 9.0 | 12.3 | 128 | 12.1 | 9.4 |
| SNCA-/- HEK293 cells | Neat | 427 | 47.5 | 11.1 | 494 | 72 | 14.6 | 120 | 29.5 | 24.6 | 461 | 25.7 | 5.6 |
| | 2 | 320 | 22.4 | 7.0 | 377 | 63.7 | 16.9 | 120 | 13.7 | 11.4 | 359 | 25.2 | 7.0 |
| | 4 | 252 | 57 | 22.6 | 182 | 15.5 | 8.5 | 93 | 2.9 | 3.1 | 279 | 14.9 | 5.3 |
| | 8 | 232 | 23.1 | 10.0 | 161 | 12.3 | 7.6 | 86 | 13.5 | 15.7 | 179 | 14.4 | 8.0 |

| | | | | | | | | | | | | | |
|---|---|---|---|---|---|---|---|---|---|---|---|---|---|
| | 16 | 140 | 11.6 | 8.3 | 136 | 9.5 | 7.0 | 84 | 6.7 | 8.0 | 171 | 8.1 | 4.7 |
| | 32 | 127 | 6.7 | 5.3 | 121 | 19.5 | 16.1 | 83 | 10.4 | 12.5 | 125 | 2.3 | 1.8 |
| | 64 | 111 | 12.1 | 10.9 | 167 | 16.4 | 9.8 | 90 | 11.7 | 13.0 | 117 | 15.6 | 13.3 |
| | 128 | 176 | 16.1 | 9.1 | 156 | 2.9 | 1.9 | 92 | 18.2 | 19.8 | 117 | 14 | 12.0 |
| | 256 | 152 | 21.5 | 14.1 | 153 | 17.1 | 11.2 | 90 | 20.1 | 22.3 | 115 | 7.6 | 6.6 |
| | 512 | 128 | 6.7 | 5.2 | 141 | 17.4 | 12.3 | 90 | 11.7 | 13.0 | 120 | 27.1 | 22.6 |
| | 1024 | 126 | 20.3 | 16.1 | 167 | 31.6 | 18.9 | 105 | 22.1 | 21.0 | 109 | 5.2 | 4.8 |
| | 2048 | 165 | 6.7 | 4.1 | 132 | 13 | 9.8 | 91 | 16 | 17.6 | 105 | 6.7 | 6.4 |
| | 4096 | 121 | 15.6 | 12.9 | 105 | 2.9 | 2.8 | 81 | 4.5 | 5.6 | 125 | 27.5 | 22.0 |
| | Blank | 126 | 16.5 | 13.1 | 123 | 6.7 | 5.4 | 96 | 22.0 | 23 | 118 | 22.6 | 19.2 |

The 2 most sensitive new total and pS129 α-synuclein assay formulations were characterized using extracts from wild-type (WT) mouse brain tissue and HEK293 cells, *Snca* knock-out mouse brain tissues, polo-like kinase 3 (PLK3)-transfected HEK293 cells, and *SNCA* knock-out HEK293 cells. Extracts were serially-diluted using assay buffer. Inter-assay variation (RSD) was defined as the standard deviation expressed as a percentage of the mean. The limit of detection (LoD) and lower limit of quantification (LLoQ) were defined as 3 and 6 standard deviations above the mean of the blank, respectively. Data represent raw assay signal measurements in arbitrary units.

**Supplementary Table 5. Evaluation of matrix effects in mouse brain tissue extracts and HEK293 cell lysates using a parallelism experiment.**

| Sample | % WT | Total α-synuclein assay | | | | pS129 α-synuclein assay | | | |
|---|---|---|---|---|---|---|---|---|---|
| | | Assay buffer | | Equivalent dilution of KO matrix | | Assay buffer | | Equivalent dilution of KO matrix | |
| | | Mean | SD | Mean | SD | Mean | SD | Mean | SD |
| 10-fold diluted WT mouse brain tissue extract | 0 | 116.0 | 2.9 | 733.0 | 75.0 | 110.0 | 11.6 | 111.0 | 7.6 |
| | 0.002 | 615.0 | 38.6 | 750.0 | 35.2 | 119.0 | 13.7 | 146.0 | 13.3 |
| | 0.005 | 2120.0 | 68.7 | 891.0 | 24.6 | 140.0 | 11.7 | 131.0 | 9.0 |
| | 0.01 | 6700.0 | 332.4 | 1878.0 | 18.5 | 158.0 | 9.0 | 140.0 | 19.7 |
| | 0.02 | 22715.0 | 1199.5 | 3747.0 | 332.0 | 167.0 | 13.0 | 140.0 | 8.1 |
| | 0.039 | 83573.0 | 1763.1 | 14633.0 | 848.6 | 204.0 | 23.2 | 142.0 | 9.5 |
| | 0.078 | 193758.0 | 4405.4 | 46588.0 | 1888.7 | 292.0 | 33.2 | 168.0 | 16.9 |
| | 0.156 | 328745.0 | 5403.2 | 108955.0 | 1519.8 | 432.0 | 36.4 | 242.0 | 23.2 |
| | 0.313 | 400612.0 | 19986.1 | 185878.0 | 17402.3 | 845.0 | 32.6 | 361.0 | 16.5 |
| | 0.625 | 430116.0 | 26996.0 | 276491.0 | 7605.2 | 2111.0 | 47.7 | 836.0 | 66.7 |
| | 1.25 | 462654.0 | 6127.1 | 315289.0 | 7690.5 | 4909.0 | 198.4 | 2120.0 | 92.2 |
| | 2.5 | 480020.0 | 16069.9 | 336507.0 | 11076.3 | 12329.0 | 341.7 | 5041.0 | 126.6 |
| | 5 | 476283.0 | 19863.4 | 324506.0 | 36664.0 | 22821.0 | 884.6 | 11551.0 | 449.6 |
| | 10 | 477771.0 | 9722.3 | 338408.0 | 10046.1 | 39129.0 | 2027.7 | 23888.0 | 302.3 |
| 100-fold diluted WT mouse brain tissue extract | 0 | 127 | 9.8 | 160 | 12.7 | 107 | 10.4 | 109 | 8.1 |
| | 0.0002 | 125 | 15.2 | 161 | 21.5 | 105 | 9.3 | 116 | 9.3 |
| | 0.0005 | 126 | 12.3 | 160 | 8.5 | 114 | 13.9 | 141 | 10.4 |
| | 0.001 | 144 | 25.1 | 216 | 35.4 | 96 | 7.1 | 117 | 12.3 |
| | 0.002 | 261 | 18.2 | 250 | 25.1 | 108 | 9 | 129 | 5.2 |
| | 0.0039 | 653 | 34.7 | 844 | 100.2 | 122 | 17.5 | 143 | 9.3 |
| | 0.0078 | 2620 | 163.4 | 2487 | 89.7 | 129 | 17.2 | 142 | 9.5 |
| | 0.0156 | 12577 | 226 | 11533 | 1315.6 | 148 | 7.5 | 148 | 9.5 |
| | 0.0313 | 51931 | 864.7 | 48368 | 3174.6 | 153 | 9.5 | 156 | 2.6 |
| | 0.0625 | 141494 | 1746.4 | 119837 | 1982.2 | 205 | 11.6 | 168 | 6.7 |
| | 0.125 | 259505 | 1965.6 | 238368 | 5006.3 | 285 | 19.3 | 274 | 14 |
| | 0.25 | 365752 | 6254.7 | 329902 | 10584.8 | 649 | 26.8 | 724 | 4.4 |
| | 0.5 | 404770 | 12546.8 | 409735 | 11360.5 | 1678 | 64.5 | 1837 | 40.5 |
| | 1 | 423557 | 6035.7 | 429215 | 9867.3 | 4655 | 194.8 | 4356 | 39.5 |
| 5-fold diluted WT HEK293 cell lysate | 0 | 115 | 12.1 | 261 | 30.5 | 101 | 13.3 | 83 | 7.1 |
| | 0.02 | 115 | 4.5 | 259 | 35.6 | 107 | 18.5 | 85 | 2.3 |
| | 0.039 | 134 | 0 | 300 | 32.4 | 118 | 16.2 | 83 | 7.1 |
| | 0.078 | 143 | 7.8 | 305 | 32.4 | 118 | 16.5 | 82 | 0.6 |
| | 0.156 | 154 | 18.2 | 360 | 65.2 | 176 | 22.7 | 113 | 22.5 |
| | 0.313 | 215 | 9.8 | 453 | 9.8 | 379 | 66.3 | 233 | 25 |
| | 0.625 | 493 | 39.9 | 794 | 73.8 | 1279 | 73.8 | 841 | 67.4 |
| | 1.25 | 1972 | 44.2 | 2496 | 81.3 | 5359 | 167.1 | 3471 | 437.6 |
| | 2.5 | 9483 | 101.7 | 8827 | 962.3 | 16345 | 1238.8 | 14320 | 497 |
| | 5 | 37689 | 890.2 | 34688 | 1512.8 | 49653 | 928.2 | 44995 | 1539.9 |
| | 10 | 117415 | 5111.7 | 98065 | 7460.8 | 128337 | 2064.5 | 108777 | 2346.6 |
| | 20 | 236526 | 5666.1 | 194142 | 3775.1 | 240675 | 3468.9 | 202285 | 2881.7 |
| 20-fold diluted WT HEK293 cell lysate | 0 | 120 | 9 | 123 | 21 | 90 | 13.3 | 87 | 16.5 |
| | 0.005 | 108 | 9.7 | 120 | 18.5 | 85 | 2.3 | 88 | 11.6 |
| | 0.01 | 112 | 30.8 | 124 | 20.3 | 85 | 7.1 | 80 | 5.2 |
| | 0.02 | 109 | 17.4 | 113 | 20.6 | 91 | 8.1 | 75 | 6.7 |
| | 0.039 | 121 | 7.2 | 139 | 34.6 | 94 | 10.4 | 82 | 12.1 |
| | 0.078 | 110 | 15.6 | 139 | 23.6 | 87 | 11 | 92 | 11.5 |
| | 0.156 | 125 | 9.3 | 143 | 25.6 | 110 | 5.2 | 83 | 16.1 |
| | 0.313 | 150 | 19.7 | 168 | 2.9 | 148 | 18.7 | 118 | 9.5 |
| | 0.625 | 222 | 34 | 252 | 11.7 | 343 | 9.8 | 270 | 33.6 |
| | 1.25 | 467 | 33.6 | 555 | 29 | 1182 | 295.8 | 1140 | 104.2 |
| | 2.5 | 1856 | 68 | 2075 | 99.8 | 4855 | 249.5 | 5233 | 280.3 |
| | 5 | 10754 | 180.9 | 9879 | 347.9 | 15518 | 420.5 | 15014 | 152.7 |

Matrix effects produced by mouse brain matrix and HEK293 cell lysate were measured for new total (#ALSU-TASYN-B, Revvity) and pS129 (#ALSU-PASYN-B, Revvity) α-synuclein assays using wild-type (WT) and α-synuclein knock-out (KO) mouse brain tissue extracts that had both been diluted 10-fold and 100-fold, as well as WT and α-synuclein KO HEK293 cell lysates that had both been diluted 5-fold and 20-fold. Data represent raw assay signal measurements in arbitrary units. SD, standard deviation.

**Supplementary Table 6. Evaluation of mouse brain tissue matrix effects in total α-synuclein assay using a spike-recovery experiment.**

| Mouse WT α-synuclein spike concentration (pg/mL) | Wild-type brain | | Knock-out brain | | Assay buffer | |
|---|---|---|---|---|---|---|
| | Measured α-synuclein concentration | % recovery | Measured α-synuclein concentration | % recovery | Measured α-synuclein concentration | % recovery |
| 10 | 2.06 | 20.63 | 4.29 | 42.89 | 4.02 | 40.23 |
| 20 | 9.58 | 47.88 | 17.84 | 89.21 | 16.40 | 81.99 |
| 40 | 39.26 | 98.15 | 39.56 | 98.89 | 38.07 | 95.16 |
| 80 | 70.41 | 88.01 | 74.15 | 92.68 | 76.78 | 95.98 |
| 160 | 138.47 | 86.54 | 157.53 | 98.46 | 164.26 | 102.66 |
| 320 | 296.74 | 92.73 | 316.50 | 98.91 | 316.03 | 98.76 |
| 640 | 775.55 | 121.18 | 668.75 | 104.49 | 894.59 | 139.78 |

Wild-type mouse α-synuclein was spiked into WT and *Snca* KO mouse brain tissue extracts diluted 2000-fold and 100-fold, respectively, as well as assay buffer (AB), to final concentrations ranging between 10-640pg/mL. Spike recovery was assessed using the new total α-synuclein assay (#ALSU-TASYN-B, Revvity) and a standard curve generated for purified mouse WT α-synuclein. Assay signal data are expressed as arbitrary units.

**Supplementary Table 7. Evaluation of mouse brain tissue matrix effects in pS129 α-synuclein assay using a spike-recovery experiment.**

| Mouse pS129 α-synuclein spike concentration (pg/mL) | Wild-type brain | | Knock-out brain | | Assay buffer | |
|---|---|---|---|---|---|---|
| | Measured α-synuclein concentration | % recovery | Measured α-synuclein concentration | % recovery | Measured α-synuclein concentration | % recovery |
| 0.625 | 0.26 | 41.60 | 0.40 | 64.37 | 0.37 | 59.77 |
| 1.25 | 1.04 | 83.50 | 1.28 | 102.47 | 1.31 | 104.49 |
| 2.5 | 1.82 | 72.99 | 2.14 | 85.53 | 2.30 | 92.18 |
| 5 | 4.04 | 80.80 | 4.16 | 83.12 | 4.39 | 87.87 |
| 10 | 9.10 | 91.05 | 9.63 | 96.27 | 9.07 | 90.71 |
| 20 | 16.77 | 83.84 | 17.72 | 88.58 | 19.24 | 96.21 |
| 40 | 44.67 | 111.68 | 42.62 | 106.55 | 44.25 | 110.63 |

Mouse pS129 α-synuclein was spiked into WT and *Snca* KO mouse brain tissue extracts diluted 2000-fold and 100-fold, respectively, as well as assay buffer (AB), to final concentrations ranging between 0.625-40pg/mL. Spike recovery was assessed using the new pS129 α-synuclein assay (#ALSU-PASYN-B, Revvity) and a standard curve generated for purified mouse pS129 α-synuclein. Assay signal data are expressed as arbitrary units.

**Supplementary Table 8. Sham and PFF mouse brain tissue extract dilutions prior to SureFire Ultra assay measurement.**

| Assay | Mouse model | Treatment | Tissue fraction | Fold dilution |
|---|---|---|---|---|
| **Total** | Wild-type | Sham | Homogenate | 1000 |
| | Wild-type | Sham | PBS | 1000 |
| | Wild-type | Sham | TrX | 500 |
| | Wild-type | Sham | SDS | 50 |
| | Wild-type | PFF | Homogenate | 1000 |
| | Wild-type | PFF | PBS | 1000 |
| | Wild-type | PFF | TrX | 500 |
| | Wild-type | PFF | SDS | 50 |
| **pS129** | Wild-type | Sham | Homogenate | 100 |
| | Wild-type | Sham | PBS | 100 |
| | Wild-type | Sham | TrX | 50 |
| | Wild-type | Sham | SDS | 50 |
| | Wild-type | PFF | Homogenate | 100 |
| | Wild-type | PFF | PBS | 100 |
| | Wild-type | PFF | TrX | 50 |
| | Wild-type | PFF | SDS | 50 |

Whole tissue homogenates, as well as PBS-soluble, TrX-soluble and SDS-soluble tissue fractions, from sham and PFF mice were pre-diluted prior to measurement of total and pS129 α-synuclein using new SureFire Ultra assays to ensure values were within the linear dynamic range of each assay.

**Supplementary Table 9. Percentage of total α-synuclein recovered in PBS, Triton and SDS brain tissue fractions from PFF and sham mice.**

| Brain region | Pathological burden | Treatment | PBS | | | TrX | | | SDS | | |
|---|---|---|---|---|---|---|---|---|---|---|---|
| | | | Mean | SD | n | Mean | SD | n | Mean | SD | n |
| MC | Severe | PFF | 93.85 | 1.47 | 9 | 5.67 | 1.47 | 9 | 0.49 | 0.17 | 9 |
| MC | Severe | Sham | 96.62 | 0.92 | 8 | 3.33 | 0.92 | 8 | 0.05 | 0.06 | 8 |
| ACC | Severe | PFF | 88.41 | 4.34 | 9 | 10.76 | 3.99 | 9 | 0.83 | 0.47 | 9 |
| ACC | Severe | Sham | 92.42 | 1.75 | 8 | 7.51 | 1.79 | 8 | 0.07 | 0.08 | 8 |
| SSC | Severe | PFF | 90.51 | 2.74 | 9 | 8.71 | 2.56 | 9 | 0.78 | 0.31 | 9 |
| SSC | Severe | Sham | 92.90 | 2.21 | 8 | 6.42 | 1.59 | 8 | 0.18 | 0.03 | 8 |
| AMG | Severe | PFF | 91.77 | 1.82 | 9 | 7.76 | 1.76 | 9 | 0.47 | 0.15 | 9 |
| AMG | Severe | Sham | 94.52 | 0.42 | 8 | 5.40 | 0.47 | 8 | 0.08 | 0.08 | 8 |
| OLF | Moderate | PFF | 91.81 | 1.38 | 9 | 8.19 | 1.38 | 9 | 0.00 | 0.00 | 9 |
| OLF | Moderate | Sham | 93.24 | 1.65 | 8 | 6.76 | 1.65 | 8 | 0.00 | 0.00 | 8 |
| STR | Moderate | PFF | 94.72 | 1.12 | 9 | 5.05 | 1.12 | 9 | 0.23 | 0.03 | 9 |
| STR | Moderate | Sham | 93.27 | 1.27 | 8 | 6.55 | 1.24 | 8 | 0.18 | 0.04 | 8 |
| HIP | Moderate | PFF | 99.24 | 0.10 | 8 | 0.33 | 0.13 | 9 | 0.46 | 0.06 | 8 |
| HIP | Moderate | Sham | 99.39 | 0.14 | 8 | 0.21 | 0.10 | 8 | 0.40 | 0.07 | 8 |
| VMB | Moderate | PFF | 99.27 | 0.48 | 9 | 0.29 | 0.14 | 9 | 0.44 | 0.36 | 9 |
| VMB | Moderate | Sham | 99.40 | 0.29 | 8 | 0.22 | 0.08 | 8 | 0.38 | 0.26 | 8 |
| DMB | No | PFF | 99.83 | 0.05 | 8 | 0.17 | 0.05 | 8 | 0.00 | 0.00 | 9 |
| DMB | No | Sham | 99.90 | 0.02 | 8 | 0.10 | 0.02 | 8 | 0.00 | 0.00 | 8 |
| CB | No | PFF | 99.79 | 0.11 | 8 | 0.21 | 0.11 | 8 | 0.00 | 0.00 | 9 |
| CB | No | Sham | 99.83 | 0.11 | 8 | 0.17 | 0.11 | 8 | 0.00 | 0.00 | 8 |

ACC, anterior cingulate cortex; AMG, amygdala; CB, cerebellum; DMB, dorso-medial midbrain; HIP, hippocampus; MC, motor cortex; OLF, olfactory bulb; SD, standard deviation; SSC, somatosensory cortex; STR, striatum; VMB, ventral midbrain.

**Supplementary Table 10. Percentage of pS129 α-synuclein recovered in PBS, Triton and SDS brain tissue fractions from PFF and sham mice.**

| Brain region | Pathological burden | Treatment | PBS | | | TrX | | | SDS | | |
|---|---|---|---|---|---|---|---|---|---|---|---|
| | | | Mean | SD | n | Mean | SD | n | Mean | SD | n |
| MC | Severe | PFF | 74.77 | 6.89 | 9 | 6.06 | 2.20 | 9 | 19.17 | 7.14 | 9 |
| MC | Severe | Sham | 99.47 | 0.17 | 8 | 0.53 | 0.17 | 8 | 0.00 | 0.00 | 8 |
| ACC | Severe | PFF | 50.88 | 15.07 | 9 | 8.63 | 2.89 | 9 | 40.48 | 13.63 | 9 |
| ACC | Severe | Sham | 99.42 | 0.32 | 8 | 0.58 | 0.32 | 8 | 0.00 | 0.00 | 8 |
| SSC | Severe | PFF | 57.12 | 11.84 | 9 | 8.51 | 3.58 | 9 | 34.37 | 10.13 | 9 |
| SSC | Severe | Sham | 99.53 | 0.10 | 8 | 0.47 | 0.10 | 8 | 0.00 | 0.00 | 8 |
| AMG | Severe | PFF | 68.93 | 9.51 | 9 | 4.91 | 2.40 | 9 | 26.16 | 9.51 | 9 |
| AMG | Severe | Sham | 99.49 | 0.14 | 8 | 0.51 | 0.14 | 8 | 0.00 | 0.00 | 8 |
| OLF | Moderate | PFF | 98.92 | 0.56 | 9 | 1.08 | 0.56 | 9 | 0.00 | 0.00 | 9 |
| OLF | Moderate | Sham | 99.73 | 0.12 | 8 | 0.27 | 0.12 | 8 | 0.00 | 0.00 | 8 |
| STR | Moderate | PFF | 79.56 | 8.58 | 9 | 1.16 | 0.48 | 9 | 19.29 | 8.62 | 9 |
| STR | Moderate | Sham | 99.06 | 0.53 | 8 | 0.94 | 0.53 | 8 | 0.00 | 0.00 | 8 |
| HIP | Moderate | PFF | 72.23 | 18.80 | 9 | 0.66 | 0.30 | 9 | 27.11 | 18.93 | 9 |
| HIP | Moderate | Sham | 99.58 | 0.19 | 8 | 0.42 | 0.19 | 8 | 0.00 | 0.00 | 8 |
| VMB | Moderate | PFF | 74.29 | 20.52 | 9 | 1.04 | 0.35 | 9 | 24.67 | 20.48 | 9 |
| VMB | Moderate | Sham | 99.69 | 0.16 | 8 | 0.31 | 0.16 | 8 | 0.00 | 0.00 | 8 |
| DMB | No | PFF | 99.38 | 0.19 | 8 | 0.57 | 0.22 | 9 | 0.00 | 0.00 | 8 |
| DMB | No | Sham | 99.74 | 0.13 | 8 | 0.26 | 0.13 | 8 | 0.00 | 0.00 | 8 |
| CB | No | PFF | 99.70 | 0.12 | 9 | 0.30 | 0.12 | 9 | 0.00 | 0.00 | 9 |
| CB | No | Sham | 99.86 | 0.10 | 8 | 0.14 | 0.10 | 8 | 0.00 | 0.00 | 8 |

ACC, anterior cingulate cortex; AMG, amygdala; CB, cerebellum; DMB, dorso-medial midbrain; HIP, hippocampus; MC, motor cortex; OLF, olfactory bulb; SD, standard deviation; SSC, somatosensory cortex; STR, striatum; VMB, ventral midbrain.

**Supplementary Table 11. Amount of α-synuclein aggregation quantified in PBS and Triton brain tissue fractions from PFF and sham mice using the LEGEND MAX α-Synuclein Aggregate ELISA.**

| Brain region | Pathological burden | Treatment | PBS | | | TrX | | |
|---|---|---|---|---|---|---|---|---|
| | | | Mean | SD | n | Mean | SD | n |
| MC | Severe | PFF | 15.96 | 8.82 | 9 | 19.06 | 10.61 | 9 |
| MC | Severe | Sham | N.D. | N.D. | 0 | N.D. | N.D. | 0 |
| ACC | Severe | PFF | 12.49 | 5.45 | 9 | 22.27 | 14.22 | 9 |
| ACC | Severe | Sham | 0.68 | N.D. | 1 | N.D. | N.D. | 0 |
| SSC | Severe | PFF | 14.44 | 4.16 | 9 | 30.14 | 16.06 | 9 |
| SSC | Severe | Sham | 0.32 | N.D. | 1 | N.D. | N.D. | 0 |
| AMG | Severe | PFF | 5.24 | 5.05 | 7 | 8.40 | 7.20 | 8 |
| AMG | Severe | Sham | 0.17 | N.D. | 1 | 2.25 | N.D. | 1 |
| OLF | Moderate | PFF | 11.27 | N.D. | 1 | 1.82 | 1.17 | 4 |
| OLF | Moderate | Sham | N.D. | N.D. | 0 | 1.09 | N.D. | 1 |
| STR | Moderate | PFF | 0.81 | N.D. | 1 | 2.88 | 3.08 | 7 |
| STR | Moderate | Sham | N.D. | N.D. | 0 | 0.28 | N.D. | 1 |
| HIP | Moderate | PFF | N.D. | N.D. | 0 | 2.71 | 3.77 | 3 |
| HIP | Moderate | Sham | N.D. | N.D. | 0 | 2.89 | N.D. | 1 |
| VMB | Moderate | PFF | 9.52 | 13.43 | 2 | 1.41 | 1.90 | 7 |
| VMB | Moderate | Sham | 0.05 | N.D. | 1 | 1.21 | N.D. | 1 |
| DMB | No | PFF | 1.23 | N.D. | 1 | N.D. | N.D. | 0 |
| DMB | No | Sham | N.D. | N.D. | 0 | N.D. | N.D. | 0 |
| CB | No | PFF | N.D. | N.D. | 0 | 0.90 | N.D. | 1 |
| CB | No | Sham | N.D. | N.D. | 0 | 0.57 | N.D. | 1 |

Mean and standard deviation (SD) values represent ng α-synuclein/mg total protein, while *n* represents the number of samples registering signals above the assay's lower limit of quantification (0.015ng/mL). Some regions had no samples with detectable aggregated α-synuclein, which were subsequently marked as not determined (N.D.). Standard deviation could not be determined in regions with less than 2 measurements. ACC, anterior cingulate cortex; AMG, amygdala; CB, cerebellum; DMB, dorso-medial midbrain; HIP, hippocampus; MC, motor cortex; OLF, olfactory bulb; SSC, somatosensory cortex; STR, striatum; VMB, ventral midbrain.
